# RBPMS and RBPMS2 Cooperate to Safeguard Cardiac Splicing

**DOI:** 10.1101/2024.11.07.622565

**Authors:** Tongbin Wu, Zeyu Chen, Zengming Zhang, Xiaohai Zhou, Yusu Gu, Frank A. Dinenno, Ju Chen

**Affiliations:** Department of Biological Sciences and Translational Medicine, Masonic Medical Research Institute, 2150 Bleecker Street, Utica, NY, 13501, USA; Department of Medicine, University of California San Diego, 9500 Gilman Drive, La Jolla, CA, 92093, USA

**Keywords:** RNA splicing, cardiomyocyte, sarcomere, splicing factors, heart development

## Abstract

**Background:** Mutations in cardiac splicing factors (SFs) cause cardiomyopathy and congenital heart disease, underscoring the critical role of SFs in cardiac development and disease. Cardiac SFs are implicated to cooperatively regulate the splicing of essential cardiac genes, but the functional importance of their collaboration remains unclear. RNA Binding Protein with Multiple Splicing (RBPMS) and RBPMS2 are SFs involved in heart development and exhibit similar splicing regulatory activities *in vitro*, but it is unknown whether they cooperate to regulate splicing *in vivo*.

**Methods:** *Rbpms* and *Rbpms2* single or double cardiomyocyte (CM)-specific knockout (KO) mice were generated and analyzed for cardiac phenotypes. RNA sequencing was performed to assess gene expression and splicing changes in single and double KOs. *In silico* analyses were used to dissect the mechanisms underlying distinct and overlapping roles of RBPMS and RBPMS2 in heart development.

**Results:** Mice lacking both RBPMS and RBPMS2 in CMs died before embryonic day 13.5 and developed sarcomere disarray, whereas *Rbpms* or *Rbpms2* single CM-specific KO mice had normal sarcomere assembly and survived to adulthood. Defective sarcomere assembly is likely owing to the widespread mis-splicing of genes essential for cardiac contraction in double KO mice, underscoring the overlapping role of RBPMS and RBPMS2 in splicing regulation. Mechanistically, we found RBPMS and RBPMS collectively promote cardiac splicing program while repressing non-cardiac splicing programs. Moreover, RNA splicing maps suggested that the binding location of RBPMS and RBPMS2 on pre-mRNA dictates whether they function as splicing activators or repressors. Lastly, the requirement for RBPMS and/or RBPMS2 for splicing regulation arises from intrinsic features of the target exons.

**Conclusions:** Our results demonstrate that RBPMS and RBPMS2 work in concert to safeguard the splicing of genes essential for cardiac contraction, highlighting the importance of SF collaboration in maintaining cardiac splicing signature, which should be taken into consideration when devising future therapeutic approaches through modulating the activity of SFs.

**Novelty and Significance:** *What Is Known?:*

- Mutations in cardiac splicing factors (SFs) cause cardiomyopathy and congenital heart disease, and the splicing of cardiac genes is regulated by multiple SFs. However, the functional importance of the collaboration among specific cardiac SFs is unknown.
- RBPMS has emerged as a cardiac SF for sarcomere genes but is not required for sarcomere assembly.
- RBPMS2 can substitute RBPMS in *in vitro* splicing assays, yet its role in mammalian cardiomyocytes (CMs) remains unclear.

*What New Information Does This Article Contribute?:*

- RBPMS and RBPMS2 have both distinct and overlapping roles in CMs.
- RBPMS and RBPMS2 collectively contribute to the maintenance of cardiac splicing program, which is essential for sarcomere assembly and embryonic survival.
- RNA splicing map of RBPMS and RBPMS2 reveals that they can function either as splicing activators or repressors, depending on their binding locations on pre-mRNA. This study provides compelling evidence of cooperation between cardiac splicing factors during heart development, which, to our knowledge, has not been demonstrated *in vivo*. *Rbpms* and *Rbpms2* CM-specific double KO mice die in utero and exhibit sarcomere disarray, whereas single KO mice survive to adulthood with normal sarcomere structure but manifest distinct cardiac phenotypes, suggesting RBPMS and RBPMS2 possess both distinct and overlapping functions in CMs. Although mis-splicing in cardiac genes can be seen in all three KOs, the splicing signature of double KO hearts drastically shifts towards non-cardiac tissues, including more prominent mis-splicing in genes related to cardiac contractile function. Our study further reveals that the splicing regulation of RBPMS and RBPMS2 has the characteristics of “positional effects”, i.e., the binding location on pre-mRNA dictates whether they function as splicing activators or repressors; and the intrinsic features of the target exon determine the requirement for one or two RBPMS proteins for splicing regulation. Our study sheds light on the functional importance of cardiac SF cooperation in maintaining cardiac splicing signature during heart development.

## Introduction

The complexity of cardiac proteome is dramatically increased by pre-mRNA alternative splicing (AS), which generates distinct protein isoforms to meet diverse needs of cardiac development and function.^1–3^ AS is tightly controlled by splicing factors (SFs) with a wide range of sequence specificities.^4^ Mutations in cardiac SFs RBM20 and RBFOX2 cause dilated cardiomyopathy and congenital heart disease, respectively,^5–8^ underscoring the critical role of SFs in cardiac development and disease. Importantly, a recent study found that the splicing of several essential cardiac genes, including *Camk2d, Ryr2, Tpm1* and *Pdlim5*, are regulated by multiple cardiac SFs.^9^ However, the functional importance of collaborations among cardiac SFs has not been examined *in vivo*.

RBPMS (<u>R</u>NA <u>B</u>inding <u>P</u>rotein with <u>M</u>ultiple <u>S</u>plicing) SFs, consisting of RBPMS and RBPMS2, are expressed in multiple cell lineages in the heart.^10–13^ RBPMS and RBPMS2 share more than 70% amino acid sequence homology^13^ with almost identical N-terminal RNA recognition motifs (RRMs),^14^ and they regulate splicing similarly in cultured smooth muscle cells,^15^ indicating there is a potential functional interplay between them. RBPMS proteins are involved in multiple aspects of RNA metabolism including RNA splicing,^15^ translocation,^16^ translation,^17^ and degradation.^18^ Importantly, RBPMS proteins emerge as pivotal regulators of cardiac development and disease, as the expression of RBPMS2 is downregulated in heart failure patients,^19^ global *Rbpms* KO mice died perinatally with cardiovascular abnormalities,^12^ and *Rbpms2*-null zebrafish displayed reduced cardiac output and myofibrillar disarray.^20^ However, the distinct and overlapping roles of RBPMS proteins in the heart, and the molecular mechanisms by which RBPMS proteins regulate the splicing of essential genes during heart development, have not been determined.

In this study, we generated *Rbpms* and *Rbpms2* floxed alleles and used them to construct cardiomyocyte (CM)-specific single and double knockout (KO) mice. Although both *Rbpms* CM-specific single KO mice (*Rbpms^cmKO^*) and *Rbpms2* CM-specific single KO mice (*Rbpms2^cmKO^*) survive to adulthood, *Rbpms/Rbpms2* CM-specific double KO mice (*Rbpms/Rbpms2^dcmKO^*) display severe cardiac defects, including sarcomere disarray, and die before embryonic day 13.5 (E13.5). These results indicate that RBPMS and RBPMS2 share a critical redundant function in CMs during heart development. To investigate the underlying molecular mechanism by which RBPMS and RBPMS2 are required for CM development, we performed RNA sequencing (RNA-seq) on single and double KOs and found RBPMS and RBPMS2 collectively regulate the splicing of sarcomere and other genes related to cardiac muscle contraction. Additionally, RBPMS proteins collaborate to maintain the cardiac splicing program, as the splicing signature of *Rbpms/Rbpms2^dcmKO^* shifts to that of non-cardiac tissues while single KOs largely retained cardiac splicing signature. Furthermore, RNA splicing maps suggested that RBPMS and RBPMS2 could function either as splicing activators or repressors, depending on their binding locations on the pre-mRNA. The requirement for RBPMS or RBPMS2 for splicing regulation arises from intrinsic features of the target exon.

Our results demonstrated that RBPMS and RBPMS2 work in concert to safeguard the splicing of genes essential for cardiac contractile function, highlighting the importance of SF collaboration in maintaining cardiac splicing signature, which should be taken into consideration when devising future therapeutic approaches that modulate the activity of SFs.

## Methods

### Mice

*Rbpms* and *Rbpms2* floxed mice were generated with CRISPR-Cas9 using a previously described method.^21^ Cardiomyocyte (CM)-specific *Rbpms* or *Rbpms2* knockout (KO) mice were created by crossing *Rbpms* or *Rbpms2* floxed mice with *Xmlc2-Cre* mice.^22^ CM-specific *Rbpms* and *Rbpms2* double KO mice were created by crossing *Rbpms* floxed mice with *Rbpms2* floxed mice, and then with *Xmlc2-Cre* mice. All mice were of C57BL/6NCrl or C57BL/6J background. Genotypes of mice were determined by polymerase chain reaction (PCR) analysis using embryonic yolk sac or tail extracts and *Rbpms^fl/fl^*primers (forward: 5’-GAATGCCTTTTCTCTATGGGAAC -3’, reverse: 5’-TGCAAACATCCCCGTTCTTC -3’), *Rbpms2^fl/fl^* primers (forward: 5’-TCCTTGCTGTCTTGCTTCTG -3’, reverse: 5’-GGTTGAAAGCTTCTGACTTCAC -3’), *Cre* primers (forward: 5’-GTTCGCAAGAACCTGATGGACA-3’, reverse: 5’-CTAGAGCCTGTTTTGCACGTTC-3’), All animal procedures were performed in accordance with the National Institutes of Health Guide for the Care and Use of Laboratory Animals and approved by the Institutional Animal Care and Use Committees of the Masonic Medical Research Institute and the University of California San Diego.

### Echocardiography

Echocardiography was performed using a VisualSonics Vevo 2100 ultrasound system (FUJIFILM) with a 32-55MHz linear transducer, as previously described.^23^ Percentage fractional shortening (%FS) was used as an indicator of cardiac systolic function. Left ventricular (LV) internal diameter at end-diastole (LVIDd) and LV internal diameter at end-systole (LVIDs) were measured on M-mode images.

### Western blots

Mouse hearts were dissected and snap-frozen in liquid nitrogen. Frozen hearts were homogenized in RIPA buffer (50 mM Tris-Cl pH 7.4, 150 mM Sodium Chloride, 1% NP-40, 0.1% SDS, 0.5% Sodium Deoxycholate, 1 mM EDTA) using a handheld pellet pestle (Sigma-Aldrich). Protein concentration was determined using Micro BCA Protein Assay Kit (Thermo Fisher Scientific). Protein samples were resolved on Bolt 4%-12% Bis-Tris gels (Life Technologies) and transferred to PVDF membranes (Bio-Rad), which were blocked with Tris Buffered Saline with 0.2% Tween-20 (TBST) supplemented with 5% BSA for 1 hour at room temperature (RT), and incubated with primary antibodies overnight at 4°C. Membranes were then washed with TBST and incubated with HRP-conjugated secondary antibodies for one hour before immunoreactive protein bands were visualized using enhanced chemiluminescence (ECL) reagent (Bio-Rad) and captured by Bio-Rad ChemiDoc Imaging System. Catalogue numbers for antibodies used in western blots in this study: pan-RBPMS, 15187-1-AP (Proteintech); pan-actin, MAB1501 (Sigma-Aldrich); α-actinin-2, A7811 (Sigma-Aldrich).

### Histology, Immunofluorescence and EdU Labelling

Histology, immunofluorescence and 5-ethynyl-2**’-**deoxyuridine (EdU) labelling were performed as previously described.^24^ Mouse hearts were dissected and fixed with 4% PFA overnight at 4°C, then incubated in 5%, 10%, 15%, 20% sucrose in PBS, embedded in OCT Tissue-Tek (Thermo Fisher Scientific), and cut to 6 μm sections using a Leica CM 3050S cryostat (Leica Microsystems). For **histology**, sections were stained with hematoxylin and eosin (H&E) using a standard protocol. Images were captured using a Hamamatsu NanoZoomer 2.0HT Slide Scanning System. For **immunofluorescence**, sections were blocked with PBST (1% BSA, 0.2% Tween-20 in PBS) for one hour, and incubated with primary antibody solution (antibodies diluted in PBST supplemented with 5% donkey serum) overnight in a humidified chamber at 4°C. Sections were then washed three times with PBST and then incubated with secondary antibody solution for 2 hours at RT. Sections were counterstained with DAPI and mounted in DAKO fluorescence mounting medium (Agilent). Images were captured using Leica SP8 confocal microscope or ECHO Revolution fluorescence microscope. For **EdU labelling**, pregnant female mice were intraperitoneally injected with EdU (5-ethynyl-20-deoxyuridine, Invitrogen) two hours prior to dissection for embryos. Embryonic hearts were processed similarly as for immunofluoscence and cut to 6 μm sections. EdU positive cells were detected with Click-iT assays (Invitrogen) using Alexa Fluor 647 azide according to the manufacturer’s instructions, which was followed by primary and secondary antibody incubation similarly as immunofluorescence procedures. Images were captured using ECHO Revolution fluorescence microscope. Sources for antibodies used in immunofluorescence in this study: pan-RBPMS, 15187-1-AP (Proteintech); Nkx2-5, sc-8697 (Santa Cruz Biotechnology); EMCN, 14-5851-82 (Invitrogen); αSMA, ab7817 (Abcam); PDGFRα, AF1062 (R&D System); α-actinin-2, ab68167 (Abcam); Myom1, B4 (DSHB).

### Polymerase chain reaction (PCR)

Total RNA was extracted from embryonic or neonatal mouse hearts using TRIzol reagent per manufacturer’s instructions (Life Technologies). cDNA was synthesized using SuperScript III reverse transcriptase (Life Technologies). Semi-quantitative PCR for splicing changes were performed on an Applied Biosystems SimpliAmp thermal cycler using DreamTaq Green PCR Master Mix (Thermo Fisher Scientific) and resolved on 2% agarose gels. Quantitative real-time PCR was performed on an Applied Biosystems QuantStudio 6 Flex Real-Time PCR System using iTaq Universal SYBR Green Supermix (Bio-Rad). *18S* rRNA was used as an internal control. Primer sequences are listed below:

*Camk2d*-F: 5’-ATTGAAGAAACCAGATGGGGTA -3’

*Camk2d* -R: 5’-TCCTCAATGGTGGTGTTTGA -3’

*Alpk3*-F: 5’-GCAGGAGCTGGTGGAGATAG-3’

*Alpk3*-R: 5’-GTGAGCTGAGCAATGATGGA-3’

*Rps24*-F: 5’-CGAAAGGAACGCAAGAACAG-3’

*Rps24*-R: 5’-CCGCAGATCTACTCCTTTGG-3’

*Svil*-EF: 5’-TGCTCAATGTGGAGAACCAG-3’ (for amplifying exons 8-10 excluded mRNA)

*Svil*-ER: 5’-AGAAGGGTACCTGGCTCGG-3’ (for amplifying exons 8-10 excluded mRNA)

*Svil*-IF: 5’-TTCTGAGCGAAGCATTTCCT-3’ (for amplifying exons 8-10 included mRNA)

*Svil*-IR: 5’-CCTGGAAGGCCAGTTGATAA-3’ (for amplifying exons 8-10 included mRNA)

*Ttn*-F: 5’-CCGCAGATCTACTCCTTTGG-3’ (work with both *Ttn*-ER and *Ttn*-IR)

*Ttn*-ER: 5’-TGCTCAATGTGGAGAACCAG-3’ (for amplifying exon 47 excluded mRNA)

*Ttn*-IR: 5’-AGAAGGGTACCTGGCTCGG-3’ (for amplifying exon 47 included mRNA)

*Scn5a*-F1: 5’-GTCTCAGCCTTACGCACCTT-3’

*Scn5a*-F2: 5’-TCCTGAGAGCTCTGAAAACGA-3’

*Scn5a*-R: 5’-CCAATGAGGGCAAAGACACT-3’ (three primers per reaction)

*Ldb3*-F: 5’-CCTATTCCCATCTCCACGAC-3’

*Ldb3*-R1: 5’-GCTAAAGGTGTCCCCGAACT-3’

*Ldb3*-R2: 5’-ATCGGGGTGTTGTACTGAGC-3’ (three primers per reaction)

*Pkm*-F1: 5’-CTCCTTCAAGTGCTGCAGTG-3’

*Pkm*-F2: 5’-CACCGTCTGCTGTTTGAAGA-3’

*Pkm*-R: 5’-TCGAGTCACGGCAATGATAG-3’ (three primers per reaction)

*18S*-F: 5’-GGAAGGGCACCACCAGGAGT-3’

*18S*-R: 5’-TGCAGCCCCGGACATCTAAG-3’

### RNA Sequencing and analysis

Embryonic or neonatal mouse hearts were homogenized in TRIzol (Invitrogen) for RNA extraction according to the manufacturer’s instructions. The concentration and quality of purified RNA was assessed by Agilent Bioanalyzer 2100. cDNA libraries were constructed using an Illumina TruSeq stranded mRNA kit according to manufacturer’s instructions. Libraries were sequenced with an Illumina NovaSeq 6000 sequencer to a sequencing depth of 80-100 million reads per sample.

Raw RNA-seq reads were quality-controlled using FastQC (http://www.bioinformatics.babrah am. ac.uk/projects/fastqc) and trimmed with Trim Galore (v0.6.7) to remove low-quality bases and adapter sequences, obtaining high-quality clean reads. Trimmed reads were aligned to the mouse genome (GRCm38, Mus musculus) using HISAT2 (version 2.2.1)^25^ and gene expression quantification were quantified using StringTie (version 2.2.1)^26^. Gene counts were obtained with featureCounts^27^ and used for differential expression analysis with DESeq2^28^. Differentially expressed genes (DEGs) were identified using the following criteria: |fold change| > 1.5 and false discovery rate (FDR)< 0.05. Benjamini-Hochberg correction for multiple testing was applied to correct p-value of each gene as FDR. Lists of downregulated DEGs and upregulated DEGs were separately examined for statistical enrichment of gene ontology (GO) terms and biological pathways in Toppgene (https://toppgene.cchmc.org).

### Alternative splicing analysis

Alternative splicing events from RNA-seq data were identified using rMATS (v4.3.0)^29^. Custom scripts were employed to filter splicing events with a read coverage greater than 5 per sample to enhance detection reliability. Alternative splicing events with FDR < 0.01 and |IncLevelDifference| > 0.1 were defined as significantly differential splicing events.

### RNA map and motif enrichment analysis

Motif enrichment analysis in regions surrounding significant alternative splicing events was performed using the findMotifsGenome.pl tool from the HOMER suite^30^ to identify potential co-regulatory splicing factors. Feature analysis of alternative splicing events was conducted using Matt (v1.3.0)^31^.

### Quantification and Statistical Analysis

For CM proliferation measurement, CM number was counted with ImageJ/Fiji’s cell count function (Nkx2-5^+^ nuclei). EdU^+^/Nkx2-5^+^ nuclei (proliferative CMs) were identified by merging EdU images (green) with Nkx2-5 images (red), and using ImageJ/Fiji’s color threshold function to select for the yellow signal generated by overlaying red Nkx2-5 and green EdU signals. Double positive nuclei were then counted with ImageJ/Fiji’s cell count function. The same settings were applied to all images for measurement. Anatomical parts of the heart (e.g., LV compact layer and Interventricular Septum (IVS) thickness) were manually defined with ImageJ/Fiji’s ROI (region of interest) function.

Data are presented as mean ± standard error of the mean (s.e.m.). Statistical analysis was performed using GraphPad Prism 9 software, with Welch’s t test or Ordinary one-way ANOVA. *P*-values less than 0.05 were considered significant and reported as *p < 0.05, **p < 0.01, ***p < 0.001, ****p<0.0001.

### Data Availability

RNA-seq data have been deposited into Gene Expression Omnibus and will be available upon publication (accession numbers: GSE280806, GSE280824). Other data and study materials are available from the corresponding author on reasonable request.

## Results

### Differential requirement of RBPMS and RBPMS2 for ventricular compaction

RBPMS and RBPMS2 share a high amino acid sequence similarity and are interchangeable in *in vitro* splicing assays.^15^ However, it remains unclear whether RBPMS and RBPMS2 display similar or distinct functions during heart development. To address this question, we constructed *Rbpms* and *Rbpms2* floxed mice with CRISPR-Cas9 (Supplemental Figure 1A-B) and validated the efficacy of *Rbpms* or *Rbpms2* floxed alleles by crossing them with germline-expressing *Sox2-Cre*^32^ (Supplemental Figure 1C-D). In line with a previous report,^12^ *Rbpms* global KO mice (*Rbpms^gKO^*) died shortly after birth and displayed cardiovascular abnormalities including noncompaction cardiomyopathy and patent ductus arteriosus (PDA) (Supplemental Figure 1E-F). In contrast, *Rbpms2* global KO mice (*Rbpms2^gKO^*) survived to adulthood without overt cardiac phenotype (data not shown), suggesting that RBPMS2 is not required for ventricular compaction, or its function can be largely compensated by RBPMS.

Utilizing a pan-RBPMS antibody, we examined the subcellular localization of RBPMS and RBPMS2 using immunofluorescence (IF) on postnatal day 0 (P0) mouse heart sections. The results showed that RBPMS and RBPMS2 were localized to nuclei of CMs, vascular smooth muscle cells (SMCs), endothelial cells (ECs) and fibroblasts (FBs), and their expression is relatively higher in CMs and SMCs (Figure 1). The specificity of the pan-RBPMS antibody was verified by the significantly reduced nuclear staining of RBPMS/RBPMS2 in *Rbpms^gKO^* samples (Figure 1). The broad expression pattern of RBPMS and RBPMS2 suggests that they may play important roles in multiple cell lineages in the heart. To study the cell autonomous role of RBPMS and RBPMS2 in CMs, we generated CM-specific *Rbpms* KO (*Rbpms^cmKO^*) and *Rbpms2* KO (*Rbpms2^cmKO^*) mice by crossing *Rbpms^fl/fl^* or *Rbpms2^fl/fl^* mice with *Xmlc2-Cre* mice, which drives Cre expression specifically in CMs as early as E7.5.^22,24,33^ Juvenile *Rbpms2^cmKO^* mice did not show any cardiac morphological defect or dysfunction (Supplemental Figure 1G-H), corroborating our findings in *Rbpms2^gKO^* mice.

**Figure 1.**
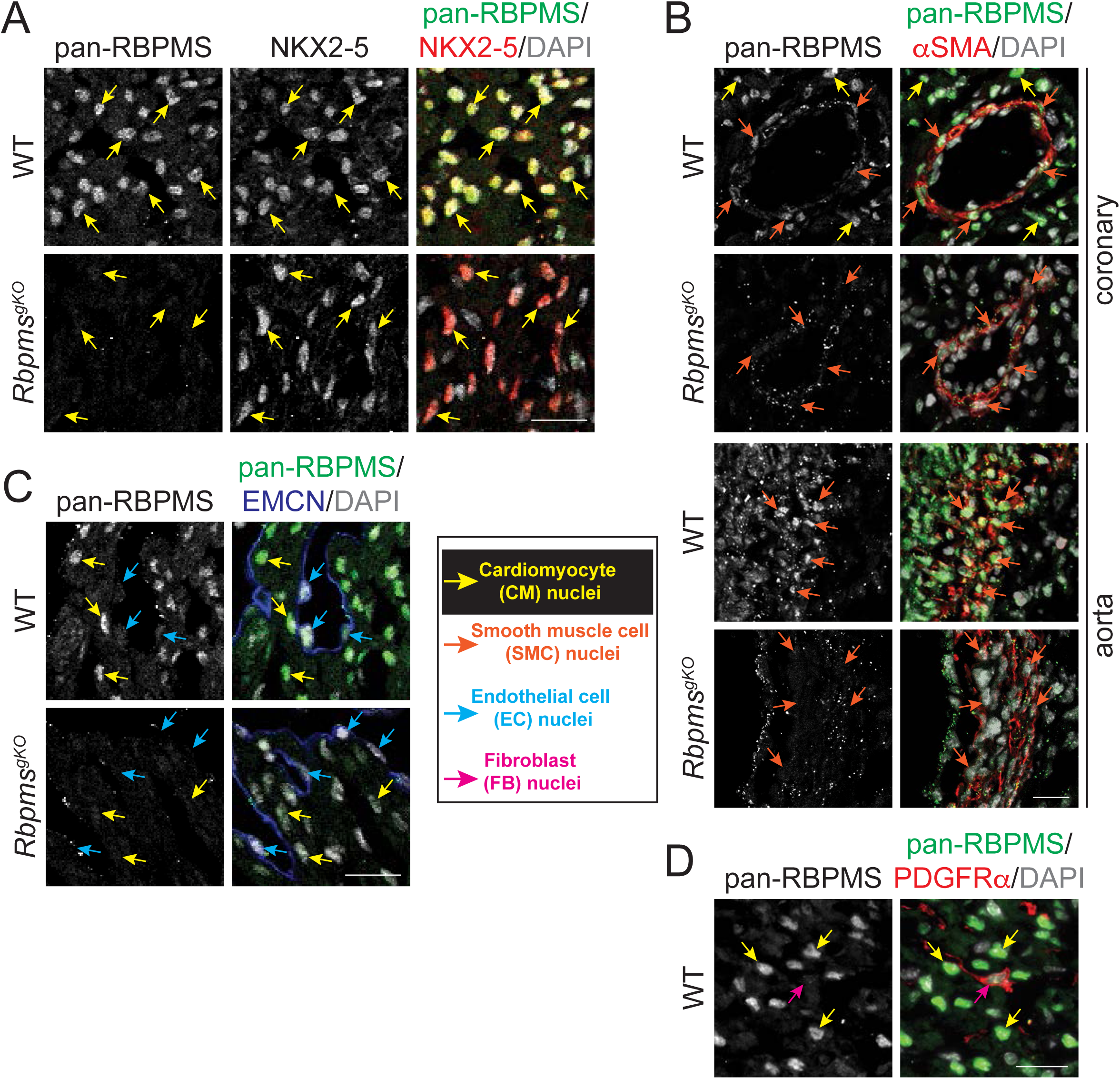
The expression and subcellular localization of RBPMS proteins in various cardiac cell types. **(A)** RBPMS and RBPMS2 (green) are localized to the nuclei (DAPI, grey) of cardiomyocytes (CMs, marked by Nkx2-5, red). **(B)** RBPMS and RBPMS2 (green) are localized to the nuclei of vascular smooth muscle cells (SMCs, marked by αSMA/α smooth muscle actin, red) in coronary vessels and aorta. **(C)** RBPMS and RBPMS2 (green) are localized to the nuclei of endothelial cells (ECs, marked by endomucin/EMCN, blue). **(D)** RBPMS and RBPMS2 (green) are localized to the nuclei of fibroblasts (FBs, marked by PDGFRα, red). Yellow arrows: CM nuclei; Orange arrows: SMC nuclei; Blue arrows: EC nuclei; Magenta arrows: FB nuclei. *Rbpms* global KO samples (*Rbpms^gKO^*) were used as negative controls. Scale bar: 20 μm.

We then sought to characterize the cardiac phenotypes of *Rbpms^cmKO^*mice. Consistent with our notion that the loss of RBPMS in multiple cell lineages in the heart contribute to the cardiac defects in *Rbpms^gKO^*mice, *Rbpms^cmKO^* mice overcame the perinatal lethality of *Rbpms^gKO^*mice and survived to adulthood (Figure 2A). Nevertheless, loss of RBPMS in CMs likely contributed to the cardiac phenotypes observed in *Rbpms^gKO^*mice because *Rbpms^cmKO^* mice also developed ventricular noncompaction (Figure 2B-C). Interestingly, prominent noncompaction was still present in one-month-old *Rbpms^cmKO^*mice (Figure 2B). In contrast, noncompaction in the recently reported *Rbpms^fl/fl^;Myh6-Cre* mice largely disappears as early as P4,^34^ probably results from the lower efficiency of *Myh6-Cre* in embryonic CMs.^35,36^ In addition, echocardiography revealed that one-month-old *Rbpms^cmKO^* mice had apparent cardiac contractile dysfunction and ventricular dilation (Figure 2D).

**Figure 2.**
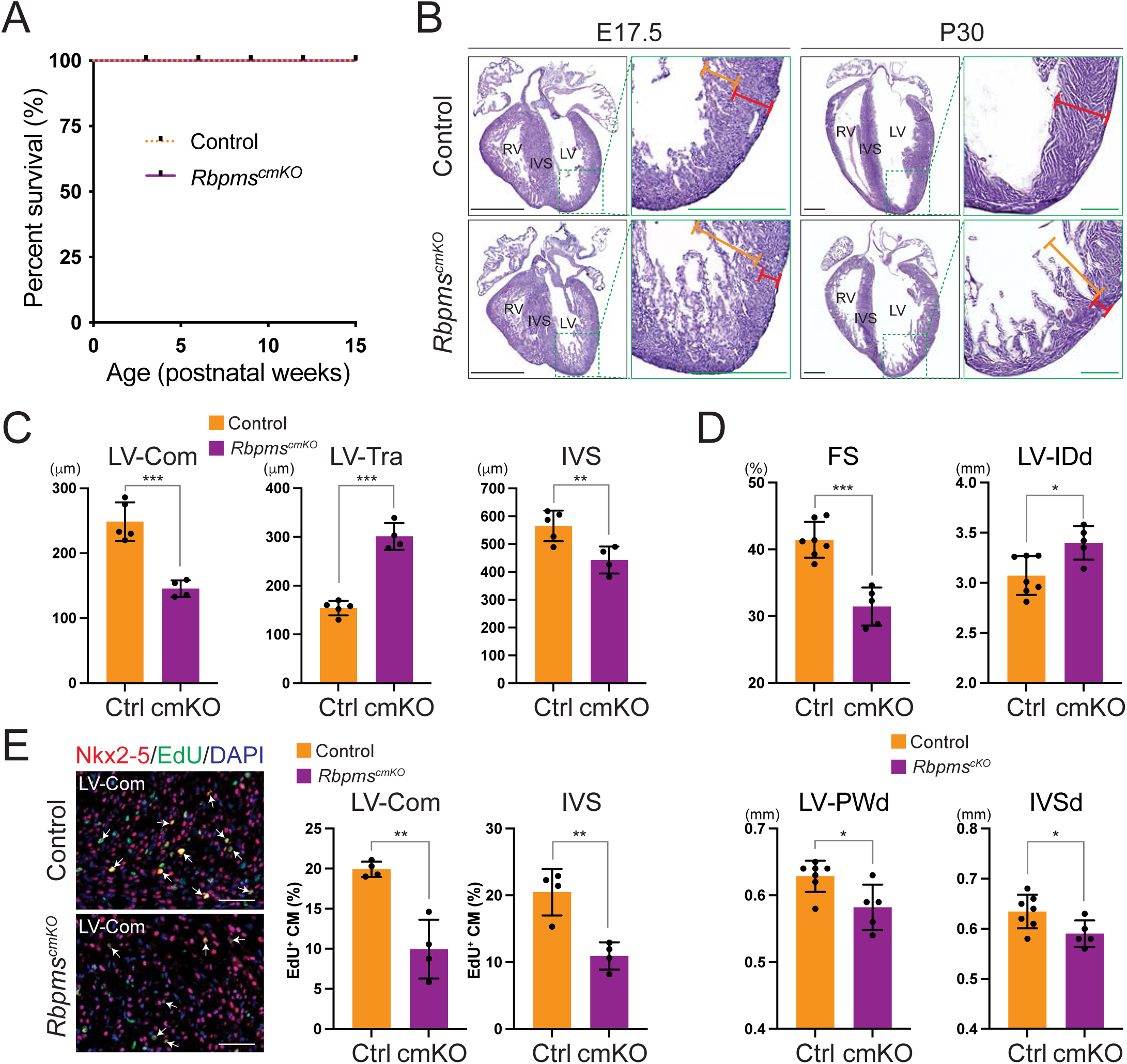
*Rbpms* CM-specific KO (*Rbpms^cmKO^*) mice develop noncompaction cardiomyopathy. **(A)** Kaplan-Meier survival curves of control (n=7) and *Rbpms^cmKO^*(n=5) mice. **(B)** Representative hematoxylin and eosin (H&E)-stained sections from control and *Rbpms^cmKO^* mice at embryonic day 17.5 (E17.5) and postnatal day 30 (P30). Orange rulers indicate left ventricle (LV) trabeculae thickness while red rulers indicate LV compact layer thickness; Scale bar: 1 mm (whole heart view); 0.5 mm (magnified view). **(C)** Quantification of the thickness of the LV compact layer (LV-Com), LV trabeculae (LV-Tra), and interventricular septum (IVS) on H&E sections from control (n=5) and *Rbpms^cmKO^* (n=4) mice at E17.5. \*\**p*<0.01, \*\*\**p*<0.001 (Welch’s *t*-test). **(D)** Echocardiography analysis of control (n=7) and *Rbpms^cmKO^* (n=5) mice at P30. \**p*<0.05, \*\*\**p*<0.001 (Welch’s *t*-test). FS: fraction shortening; LV-IDd/s: LV internal dimension, diastolic/systolic; LV-PWd: LV posterior wall thickness, diastolic; IVSd: interventricular septum thickness, diastolic. **(E) Left panel:** representative immunostaining images of LV-com of E17.5 control and *Rbpms^cmKO^* mice using antibodies against Nkx2-5 (red), EdU signal (green), and DAPI (blue). Arrows indicate EdU-positive cardiomyocytes (CMs). Scale bar: 50 μm. **Right panel:** quantification (%) of EdU-positive CMs in E17.5 control (n=4) vs. *Rbpms^cmKO^*(n=4) heart areas as indicated. \*\**p*<0.01 (Welch’s t-test).

To study the cellular basis of the noncompaction phenotype in *Rbpms^cmKO^* mice, we performed (EdU) labeling and found that CM proliferation was significantly reduced in the LV compact myocardium and IVS of embryonic day 17.5 (E17.5) *Rbpms^cmKO^*hearts (Figure 2E), which may account for the reduced thickness of these regions.

In summary, our findings indicate that RBPMS is required in CMs for ventricular compaction, and it likely functions in various cardiac cell types, as *Rbpms^cmKO^* mice only partially recapitulated the cardiovascular phenotypes of *Rbpms^gKO^* mice. On the contrary, ablating RBPMS2 globally or specifically in CMs did not affect normal heart development, suggesting the function of RBPMS2 during ventricular compaction can be largely compensated by RBPMS but not vice versa.

### RBPMS and RBPMS2 collaborate to regulate sarcomere assembly

A study in zebrafish found that RBPMS2 regulates sarcomere assembly.^20^ However, RBPMS seems to be dispensable for maintaining sarcomere structure in mice.^12,34^ One caveat is that only RBPMS2 is expressed in zebrafish heart^20^ whereas both RBPMS and RBPMS2 are expressed in murine heart (Supplemental Figure 1C-D). Thus, RBPMS and RBPMS2 may substitute each other in mammals to ensure proper sarcomere assembly. To test this possibility, we generated CM-specific *Rbpms* and *Rbpms2* double KO mice (*Rbpms/Rbpms2^dcmKO^*) to investigate overlapping functions of RBPMS proteins in CMs. Strikingly, *Rbpms/Rbpms2^dcmKO^* mice showed cardiac defects including a thinner compact layer, fewer trabeculae, and underdeveloped septum at E12.5, and died before E13.5 (Figure 3A-B and Supplemental Figure 2), which is in stark contrast with *Rbpms^cmKO^*and *Rbpms2^cmKO^* mice that survived to adulthood. These observations suggest that RBPMS and RBPMS2 share a critical redundant role which is indispensable for cardiac development.

**Figure 3.**
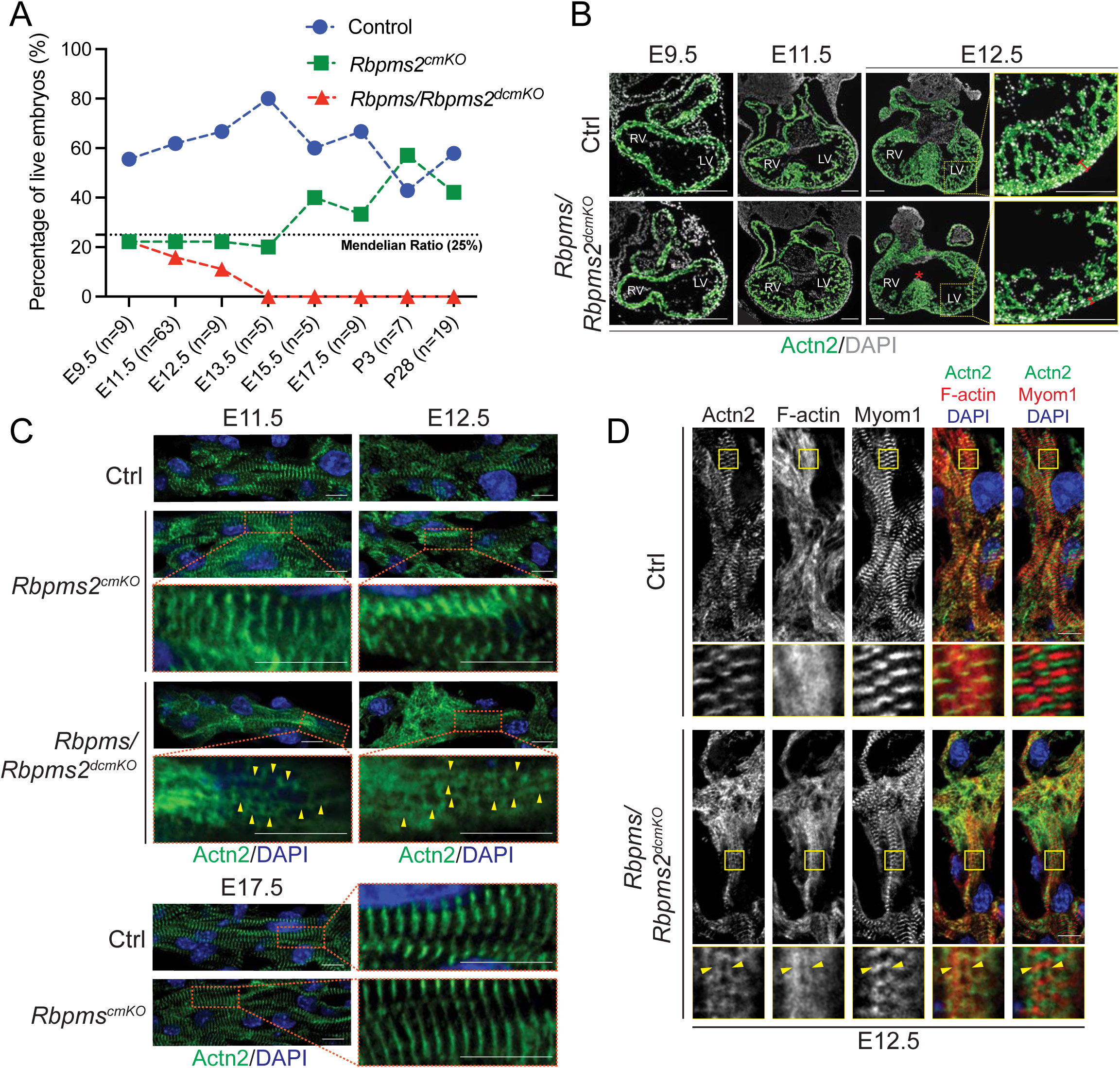
*Rbpms/Rbpms2* CM-specific double KO (*Rbpms/Rbpms2^dcmKO^*) mice died before E13.5 with sarcomere disarray. **(A)** Percentage of live embryos/pups from timed pregnancy of crossing *Rbpms^fl/fl^;Rbpms2^fl/fl^* mice with *Rbpms^fl/+^;Rbpms2^fl/fl^;Xmlc2-Cre^+^* mice. The expected Mendelian ratio of double KO is 25%. **(B)** *Rbpms/Rbpms2^dcmKO^*hearts showed morphological abnormalities starting from E12.5. Asterisk indicates underdeveloped septum. Red rulers indicate compact layer thickness. **(C-D)** Immunofluorescent images from control, *Rbpms^cmKO^, Rbpms2^cmKO^* and *Rbpms/Rbpms2^dcmKO^* at various developmental stages. Yellow arrows indicate abnormal actin bundles (AABs). Scale bar: 0.2 mm (B); 10 μm (C-D).

To determine whether RBPMS and RBPMS2 are required for sarcomere assembly, we performed immunofluorescent staining using an antibody against Z-line protein α-actinin-2 (Actn2) and observed obvious sarcomere disarray in the CMs of *Rbpms/Rbpms2^dcmKO^*(Figure 3C). In particular, *Rbpms/Rbpms2^dcmKO^* CMs formed α-actinin-2 (Actn2)-positive structures interconnecting neighboring Z-lines, which were not found in E11.5-E12.5 control or *Rbpms2^cmKO^* CMs (Figure 3C). Notably, we did not observe such structures or any abnormalities in the sarcomeres of *Rbpms^cmKO^*CMs at E17.5 (Figure 3C), a stage when *Rbpms^cmKO^* mice have already displayed cardiac phenotypes (Figure 2). We further investigated the components of the atypical structures in *Rbpms/Rbpms2^dcmKO^* mice by performing Immunofluorescent staining with phalloidin (F-actin) and myomesin 1 (Myom1, M-line marker), which revealed that these structures contain Actn2 but not Myom1 (Figure 3D), reminiscent of the “abnormal actin bundles (AABs)” found in HSPB7 KO mice.^37^ These results suggest that RBPMS and RBPMS2 cooperatively regulate sarcomere assembly in CMs.

### Modest gene expression changes consequent to loss of RBPMS and/or RBPMS2 in CMs

To determine whether global gene expression changes could account for observed cardiac phenotypes, we performed RNA sequencing (RNA-seq) on *Rbpms^cmKO^, Rbpms2^cmKO^*, and *Rbpms/Rbpms2^dcmKO^* hearts at stages prior to overt cardiac defects (*Rbpms2^cmKO^* and *Rbpms/Rbpms2^dcmKO^* at E11.5) or only with mild phenotypes (*Rbpms^cmKO^* at P0). Principal component analysis (PCA) indicated that transcriptomes of *Rbpms^cmKO^* and *Rbpms/Rbpms2^dcmKO^* underwent significant changes compared to controls while that of *Rbpms2^cmKO^* was only modestly altered (Figure 4A). We then sought to identify unique and common differentially expressed genes (DEGs) among *Rbpms^cmKO^, Rbpms2^cmKO^*, and *Rbpms/Rbpms2^dcmKO^*hearts, but only found a small fraction of DEGs that were shared among different KOs (Figure 4B).

**Figure 4.**
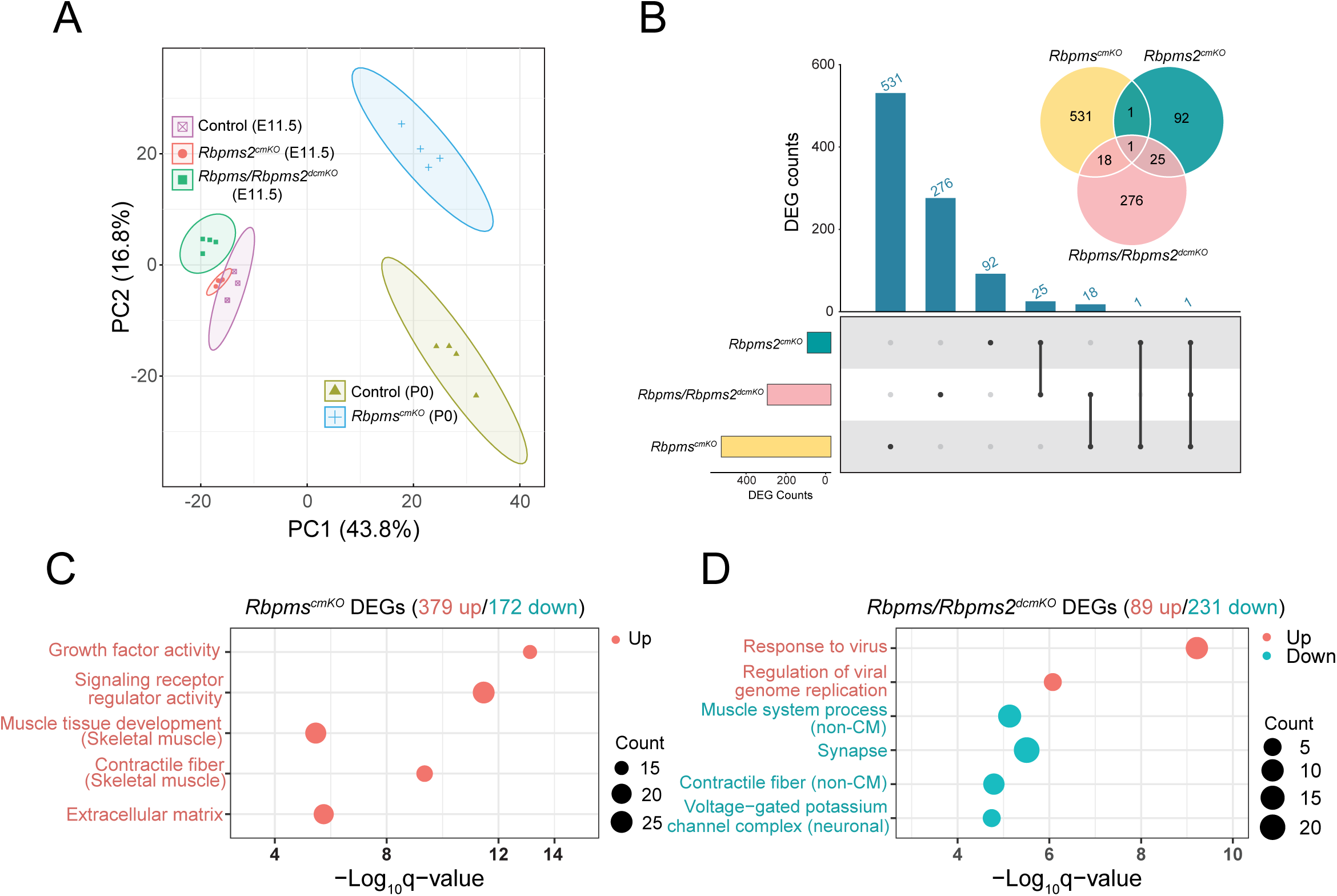
Global gene expression changes consequent to loss of RBPMS proteins individually or altogether. **(A)** Principal component analysis of RNA-seq datasets of *Rbpms^cmKO^, Rbpms2^cmKO^*, and *Rbpms/Rbpms2^dcmKO^* hearts (n=3-4). Note that *Rbpms^cmKO^* and *Rbpms/Rbpms2^dcmKO^*samples were clearly separated from their control samples whereas *Rbpms2^cmKO^*samples were not. **(B)** Overlap of differentially expressed genes (DEGs) identified in *Rbpms^cmKO^, Rbpms2^cmKO^*, and *Rbpms/Rbpms2^dcmKO^* hearts. **(C-D)** Gene ontology (GO) analysis of DEGs identified in *Rbpms^cmKO^* **(C)** and *Rbpms/Rbpms2^dcmKO^* **(D)** hearts. No GO terms were significantly enriched (Bonferroni adjusted p-value <0.01) in the DEGs of *Rbpms2^cmKO^*or downregulated DEGs in *Rbpms^cmKO^* hearts.

Gene ontology (GO) analysis revealed that upregulated DEGs in *Rbpms^cmKO^*mice enriched cardiac fibrotic and remodeling gene programs, including *Col22a1, Ctgf/Ccn2, Nppa, Nppb* (Figure 4C and Supplemental Table 1), which is in line with the diseased state of *Rbpms^cmKO^* heart. Although GO terms “muscle tissue development” and “contractile fiber” were also enriched in upregulated DEGs, they mainly consist of genes encoding skeletal muscle form of sarcomeric or calcium handling proteins (*Tpm2, Myh1, Myh2, Myh13, Myl11, Ryr1*), and cytoskeletal proteins tend to be upregulated in cardiomyopathy (*Xirp2, Ankrd1*)^38,39^ (Figure 4C and Supplemental Table 1). We also found that *p21/Cdkn1a*, a cell cycle inhibitor that represses CM proliferation in the developing heart,^40^ was upregulated in *Rbpms^cmKO^* mice (Supplemental Table 1), which may partially account for the reduced CM proliferation in *Rbpms^cmKO^*mice (Figure 2D). In contrast, downregulated DEGs in *Rbpms^cmKO^* hearts and all DEGs in *Rbpms2^cmKO^* hearts did not enrich any GO terms, suggesting that RBPMS and RBPMS2 are not required for transcription or mRNA stability of genes involved in essential biological processes during heart development.

Furthermore, none of the enriched GO term DEGs in *Rbpms/Rbpms2^dcmKO^*hearts are related to heart development and function (Figure 4D). Notably, “muscle system process” and “contractile fiber” GO terms resulted from downregulation of genes encoding non-CM form of sarcomeric and sodium/calcium handling proteins (*Myh3, Myl1, Acta1, Acta2, Myl9, Scn1a, Casq1*) (Supplemental Table 1). In summary, our findings suggest that the cardiac phenotypes in *Rbpms^cmKO^* and *Rbpms/Rbpms2^dcmKO^* hearts are unlikely caused by changes in the overall expression level of essential genes.

### RBPMS and RBPMS2 cooperate in cardiac splicing

Previous studies have demonstrated that RBPMS and RBPMS2 are *bona fide* splicing factors.^15,20^ To determine the role of RBPMS and RBPMS2 in cardiac splicing regulation, we analyzed RNA-seq data with replicate multivariate analysis of transcript splicing (rMATS)^29^ to identify dysregulated alternative splicing (AS) events in *Rbpms^cmKO^, Rbpms2^cmKO^*, and *Rbpms/Rbpms2^dcmKO^* hearts. The results showed that loss of RBPMS in CMs (*Rbpms^cmKO^*) led to significant changes in 1,012 AS events, loss of RBPMS2 in CMs (*Rbpms2^cmKO^*) led to 624 mis-spliced AS events, and loss of RBPMS and RBPMS2 altogether in CMs (*Rbpms/Rbpms2^dcmKO^*) resulted in 1,053 dysregulated AS events (ΔPercent Spliced In (ΔPSI) > 10%, False Discovery Rate (FDR) < 0.01). We focused on skipped exon (SE) and mutual exclusive exon (MXE) types which made up most of the AS events (Fig. 5A).

**Figure 5.**
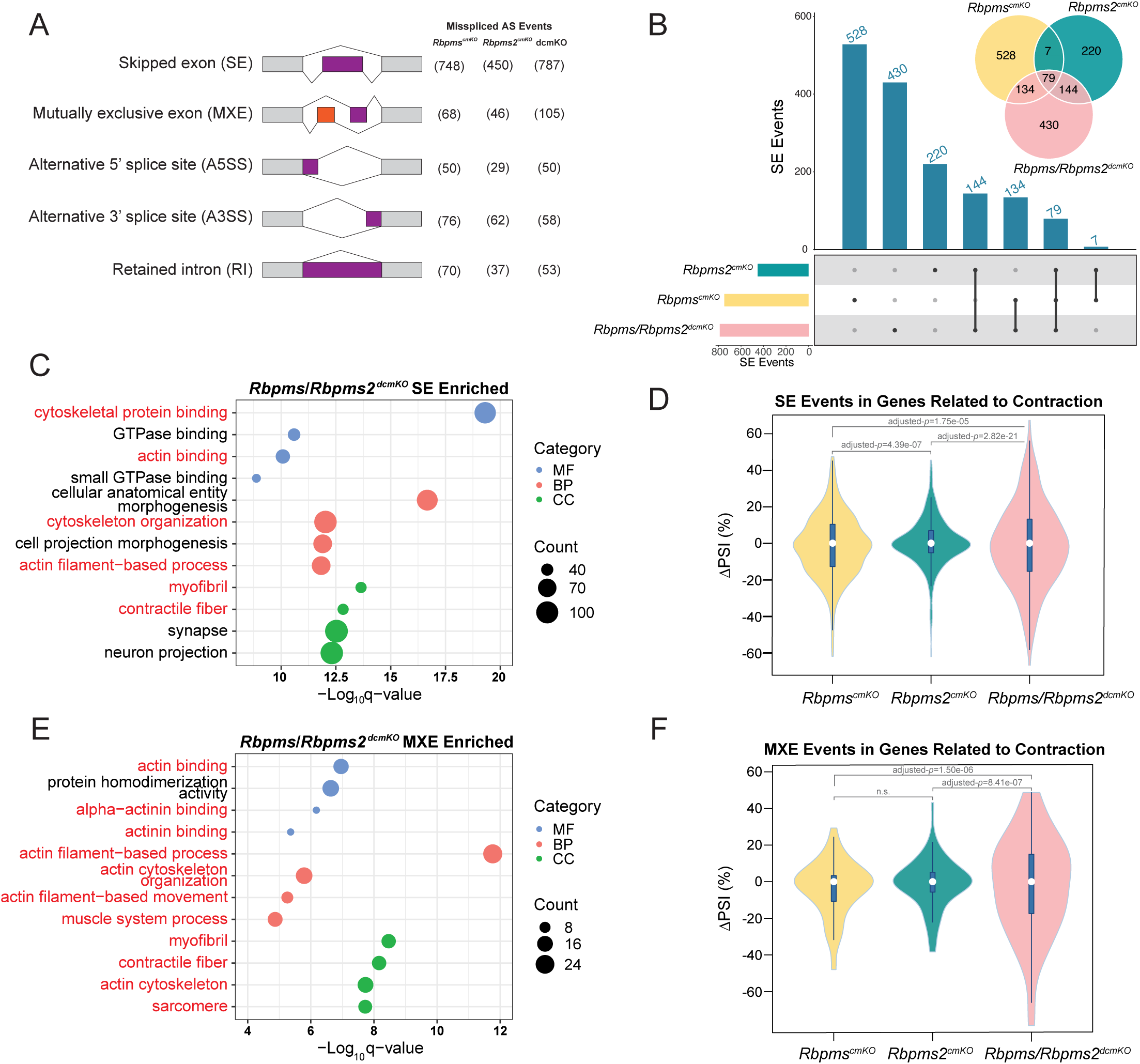
RBPMS and RBPMS2 cooperate in cardiac splicing. **(A)** Dysregulated alternative splicing (AS) event types in *Rbpms^cmKO^, Rbpms2^cmKO^*, and *Rbpms/Rbpms2^dcmKO^* hearts. Numbers of each AS type are indicated. **(B)** Overlap of skipped exon (SE) events among *Rbpms^cmKO^, Rbpms2^cmKO^*, and *Rbpms/Rbpms2^dcmKO^* hearts. **(C)** Gene ontology (GO) analysis of mis-spliced SE events in *Rbpms/Rbpms2^dcmKO^* hearts. GO terms related to cardiac contraction are highlighted in red. BP, biological process; CC, cellular component; MF, molecular function. **(D)** Violin plot showing splicing changes of SE events in genes related to cardiac contraction function (compiled from GO terms highlighted in red in Fig. 5C and Supplemental Fig. 3A; gene lists of GO terms shown in Supplemental Table 2) among *Rbpms^cmKO^, Rbpms2^cmKO^*, and *Rbpms/Rbpms2^dcmKO^* hearts. PSI, percentage spliced in. Statistical significance of differences in PSI distribution among three groups is determined by Dunn test. **(E)** Gene ontology (GO) analysis of mis-spliced mutual exclusive exon (MXE) events in *Rbpms/Rbpms2^dcmKO^* hearts. GO terms related to cardiac contraction are highlighted in red. **(F)** Violin plot illustrating splicing changes of MXE events in genes related to cardiac contraction function (compiled from GO terms highlighted in red in Fig. 5E and Supplemental Fig. 3B; gene lists of GO terms shown in Supplemental Table 2) among *Rbpms^cmKO^, Rbpms2^cmKO^*, and *Rbpms/Rbpms2^dcmKO^* hearts. Statistical significance of differences in PSI distribution among three groups is determined by Dunn test. n.s., not significant.

To identify SEs individually or jointly regulated by RBPMS and RBPMS2, we compared SEs dysregulated in *Rbpms^cmKO^, Rbpms2^cmKO^*, and *Rbpms/Rbpms2^dcmKO^*. The results indicated RBPMS and RBPMS2 regulate distinct sets of SEs in CMs (Figure 5B). Despite the overlap between SEs of *Rbpms^cmKO^* and *Rbpms2^cmKO^* being statistically significant (p-value<2.2e-16, odds ratio=4.79, Fisher’s exact test), SEs jointly regulated by RBPMS and RBPMS2 may still be undercounted due to their potential functional redundancy in CMs. Therefore, we focused on SEs that were altered in *Rbpms/Rbpms2^dcmKO^*, as these are more likely to be commonly regulated by RBPMS and RBPMS2. We found GO terms like “actin binding”, “actin filament-based process”, and “myofibril” were among the most enriched (Figure 5C and Supplemental Table 2), consistent with the sarcomere disarray phenotype observed in *Rbpms/Rbpms2^dcmKO^* hearts. Although dysregulated SE events in *Rbpms^cmKO^* or *Rbpms2^cmKO^* also enriched similar GO terms (Supplemental Figure 3A and Supplemental Table 2), *Rbpms/Rbpms2^dcmKO^* had more prominent changes in cardiac contractile function-related SEs compared to single KOs (Figure 5D), suggesting these SEs are co-regulated by RBPMS and RBPMS2. Next, to determine whether RBPMS and RBPMS2 also collaborate in controlling MXE events in CMs, we compared dysregulated MXEs among *Rbpms^cmKO^, Rbpms2^cmKO^*, and *Rbpms/Rbpms2^dcmKO^* (Supplemental Figure 3C). Despite enriching similar GO terms as *Rbpms^cmKO^* and *Rbpms2^cmKO^* (Figure 5E and Supplemental Figure 3B), MXEs in *Rbpms/Rbpms2^dcmKO^*also had more pronounced splicing changes (Figure 5F). We validated several SE and MXE events in key cardiac genes by reverse transcription PCR (RT-PCR), including *Camk2d, Alpk3, Rps24, Svil, Ttn, Scn5a, Ldb3*, and *Pkm*, all of which exhibited more dramatic splicing changes in *Rbpms/Rbpms2^dcmKO^* compared with single KOs (Supplemental Figure 4A-H).

Taken together, our findings suggest that RBPMS and RBPMS2 collaborate in maintaining proper splicing of sarcomere and other muscle contraction-related genes in CM. The redundant function of RBPMS and RBPMS2 in splicing regulation, a form of genetic compensation,^41^ ensures normal sarcomere assembly and embryonic survival in the event of genetic perturbations to either RBPMS or RBPMS2.

### The shift from cardiac to non-cardiac splicing signatures in *Rbpms/Rbpms2^dcmKO^* hearts

Close examination of AS events collectively regulated by RBPMS and RBPMS2 revealed that their splicing pattern had the tendency to shift from cardiac towards non-cardiac tissues in *Rbpms/Rbpms2^dcmKO^*. For example, exon 6 of *Rps24*, a congenital heart disease (CHD) gene,^42^ is predominantly included in the heart.^43^ However, the inclusion of exon 6 was dramatically decreased in the heart of *Rbpms/Rbpms2^dcmKO^* mice (Supplemental Figure 4C). Similarly, the muscle-specific long isoform of *Svil*, encoding Archvillin,^44^ was significantly decreased in *Rbpms/Rbpms2^dcmKO^*mice (Supplemental Figure 4D). Moreover, the cardiac isoform of *Ldb3* (also known as *Cypher*)^45^ was reduced in *Rbpms^cmKO^* mice and reduced further in *Rbpms/Rbpms2^dcmKO^* mice, while the skeletal muscle isoform of *Ldb3* was concurrently increased (Supplemental Figure 4G). These findings suggest that these cardiac-specific exons are collectively promoted by RBPMS and RBPMS2 to retain cardiac splicing patterns in CMs.

To assess whether this shift from cardiac to non-cardiac splicing patterns occurs across the genome, we first calculated PSI-based cardiac and non-cardiac splicing signatures using RNA-seq datasets obtained from various wild-type mouse tissues at E11.5 and P0,^46,47^ and visualized them in a PCA plot (Figure 6). While Dimension 2 (Dim2) mainly reflected developmental differences in AS programs among embryonic (E11.5), neonatal (P0) and adult (P63) heart samples, Dimension 1 (Dim1) clearly separated cardiac from non-cardiac tissues, consistent with the previous observation that heart has a distinct AS program.^47^ To determine the overall impact of losing RBPMS and/or RBPMS2 on cardiac AS program, we analyzed RNA-seq data of *Rbpms^cmKO^, Rbpms2^cmKO^*, and *Rbpms/Rbpms2^dcmKO^* and found heart samples from single KOs (*Rbpms^cmKO^* at E11.5 and P0; *Rbpms2^cmKO^*at E11.5) stayed at the “cardiac side” (Figure 6), whereas heart samples from *Rbpms/Rbpms2^dcmKO^* (at E11.5) moved closer to non-cardiac samples (Figure 6), suggesting that ablating RBPMS and RBPMS2 altogether dramatically shifts the cardiac splicing program towards non-cardiac splicing programs. In addition, *Rbpms/Rbpms2^dcmKO^*hearts also have tendency to lose fetal heart splicing signature as they move towards neonatal and adult heart samples (“adult heart side”, Figure 6). Collectively, our data indicate that RBPMS and RBPMS2 work in concert to maintain the unique cardiac AS signature which is required for normal heart development.

**Figure 6.**
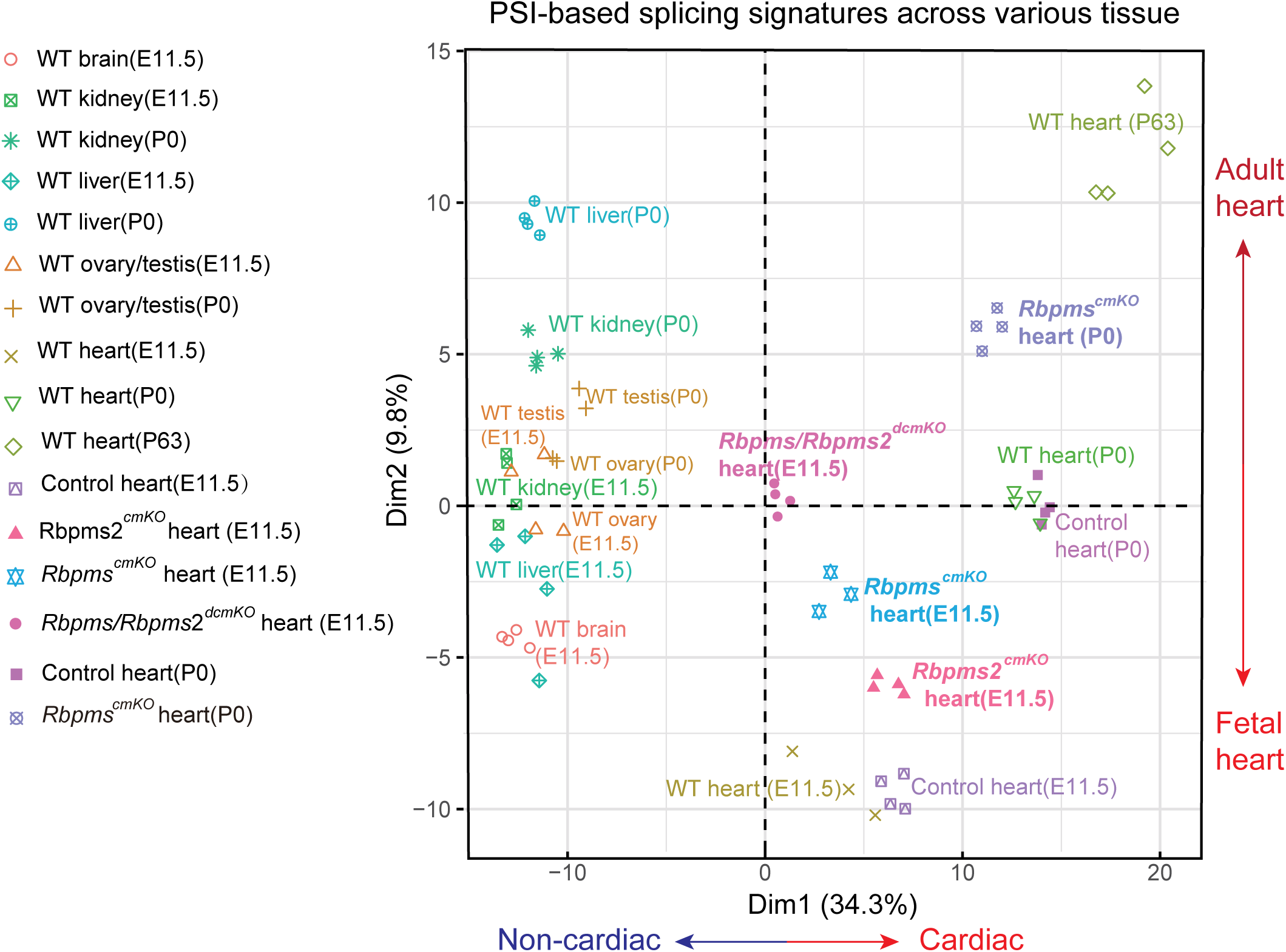
The shift from cardiac to non-cardiac splicing signatures in *Rbpms/Rbpms2^dcmKO^* hearts. Principal component analysis (PCA) plot of PSI-based splicing signatures calculated from RNA-seq datasets obtained from various wild-type (WT) mouse tissues at different developmental time points (Cardoso-Moreira et al., 2019) and RNA-seq datasets from *Rbpms^cmKO^, Rbpms2^cmKO^, Rbpms/Rbpms2^dcmKO^* and littermate control hearts at developmental stages as indicated.

### RNA maps of RBPMS and RBPMS2 in CMs

The location of SF binding sites on the pre-mRNA tends to determine the splicing outcome of their target genes, a phenomenon known as “positional effects” and is often depicted in “RNA maps”.^4,48–51^ As consensus binding motifs of SFs correspond very well with their *in vivo* binding sites,^15,48,52–54^ we used the well-defined RBPMS/RBPMS2 motif, CAC motifs separated by spacers of variable length (CACN_1-12_CAC),^14,15^ to predict the binding sites of RBPMS and RBPMS2, and further construct their RNA maps in conjunction with RNA-seq data. Specifically, we calculated the enrichment of CAC motifs around skipped exons activated or repressed only by RBPMS (splicing altered in *Rbpms^cmKO^*and *Rbpms/Rbpms2^dcmKO^* but not in *Rbpms2^cmKO^*, 662 SE events, see Figure 5B); only by RBPMS2 (splicing altered in *Rbpms2^cmKO^*and *Rbpms/Rbpms2^dcmKO^* but not in *Rbpms^cmKO^*, 364 SE events); or by both RBPMS and RBPMS2 (exon inclusion altered in all three KOs, or only in *Rbpms/Rbpms2^dcmKO^* due to functional compensation by RBPMS or RBPMS2, 509 SE events). Our analysis revealed that exons whose inclusion is promoted by RBPMS and/or RBPMS2 had strong motif enrichment proximal to the alternative 5’ splice site (ss) (blue box, Figure 7A), while exons whose inclusion is repressed by RBPMS and/or RBPMS2 had significant motif enrichment within the exon and close to the constitutive 5’ss (red boxes, Figure 7A), a pattern resembling that of neuronal splicing factor Nova^48,54^.

**Figure 7.**
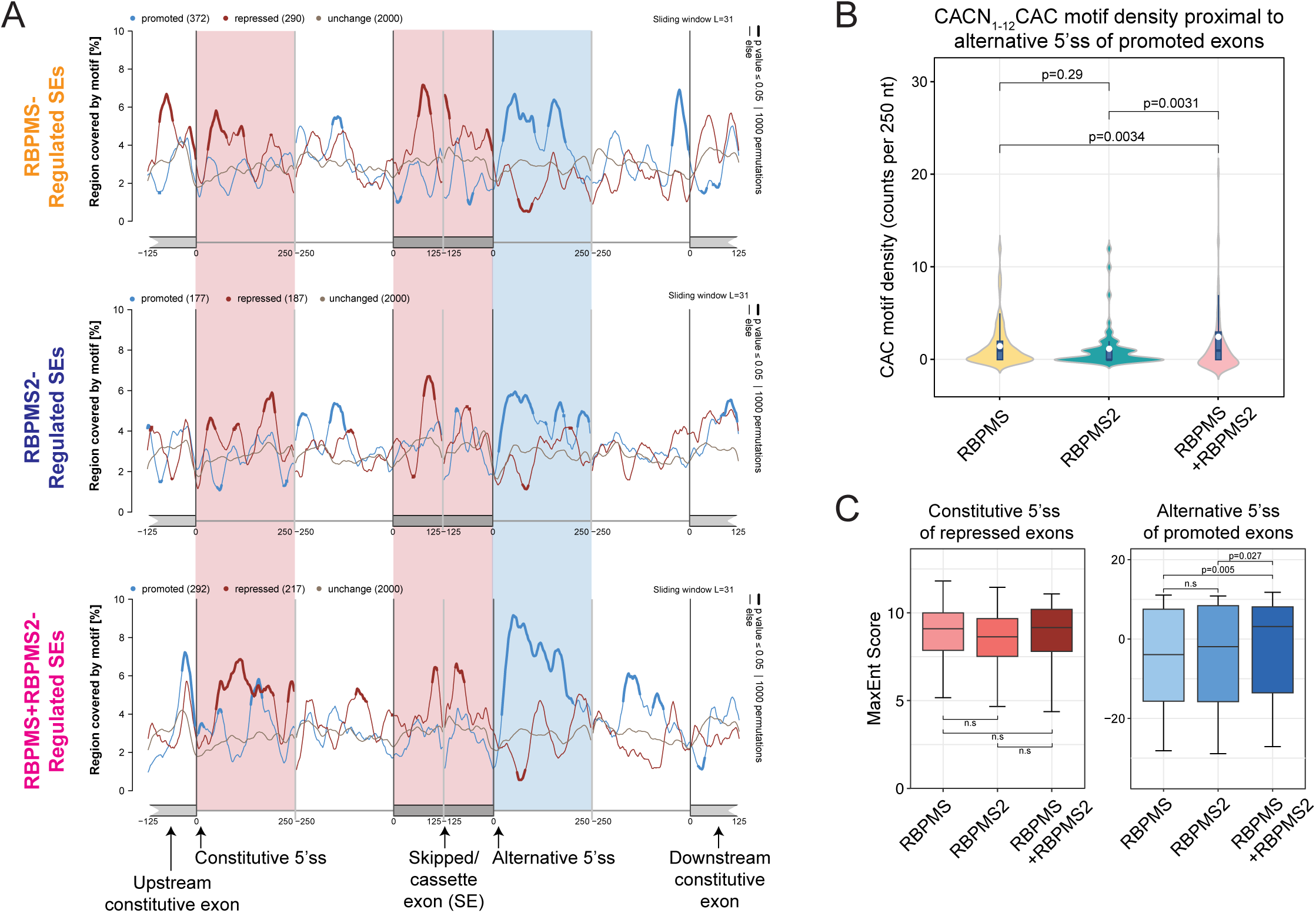
RNA maps of RBPMS and RBPMS2 in CMs. **(A)** RBPMS and RBPMS2 RNA map calculated from skipped exons (SE) regulated by RBPMS only, by RBPMS2 only or by both RBPMS and RBPMS2. Motif coverage is shown around exons promoted by RBPMS/RBPMS2 (blue lines), exons repressed by RBPMS/RBPMS2 (red lines), and exons not regulated by RBPMS/RBPMS2 (unregulated, gray lines). Statistical significance (*p*<0.05, 1000 permutations) is indicated by thicker lines. Red boxes and blue boxes indicate regions with most significant motif coverage associated with repressed (red boxes) and promoted exons (blue boxes), respectively. **(B)** Comparison of RBPMS/RBPMS2 motif coverage within blue boxes in (A) among three types of SE exons. Statistical significance was determined by pairwise Wilcoxon rank-sum tests. Benjamini-Hochberg method was applied to the obtained adjusted p-values due to multiple comparisons. n.s., not significant. **(C)** 5’ splice site (5’ss) strength at constitute 5’ss or alternative 5’ss of SE exons individually or cooperatively regulated by RBPMS and RBPMS2. Statistical significance was determined by pairwise Wilcoxon rank-sum tests. Benjamini-Hochberg method was applied to the obtained adjusted p-values due to multiple comparisons. n.s., not significant.

Because positional effects of RBPMS and RBPMS2 are similar (Figure 7A), it remains unclear what determines the requirement for RBPMS or RBPMS2 for splicing regulation. We noticed the motif enrichment appeared to be higher at exons co-promoted by RBPMS and RBPMS2 (see blue box of Figure 7A). To confirm this observation, we compared CAC motif enrichment at alternative 5’ss of exons individually or collectively promoted by RBPMS and RBPMS2 and found that exons coregulated by both RBPMS and RBPMS2 had higher CAC motif content (Figure 7B). On the other hand, we did not find a significant difference in CAC motif enrichment at constitutive 5’ss and exon bodies of exons repressed by RBPMS and/or RBPMS2 (Supplemental Figure 5A-B). These findings suggest that long stretches of CAC motifs at alternative 5’ss of skipped exons may require the binding of both RBPMS and RBPMS2 for splicing regulation.

The relative strength of splice sites has profound effects on splicing outcomes.^49,50,55^ We then asked whether the strength of 5’ss could determine the requirement for one or two RBPMS proteins for splicing regulation. Using a maximum-entropy model,^56^ we calculated the strength of constitutive 5’ss of repressed exons (left red box, Figure 7A) and alternative 5’ss of promoted exons (blue box, Figure 7A). Although there is no significant difference in constitutive 5’ss strength among exons repressed by RBPMS and/or RBPMS2 (left panel, Figure 7C), alternative 5’ss of exons promoted by both RBPMS and RBPMS2 are stronger than the ones that only need RBPMS or RBPMS2 for regulation (right panel, Figure 7C).

Taken together, our data indicate that RBPMS and RBPMS2 have positional effects in their splicing regulation, and the intrinsic features of the target exon likely determine the requirement for one or two RBPMS proteins for splicing regulation.

## Discussion

In this study, we aimed to determine the functional importance of the collaboration between RBPMS and RBPMS2 during heart development. To that end, we constructed *Rbpms* and *Rbpms2* floxed mice and used them to generate mouse models lacking one or both of RBPMS proteins specifically in CMs. Surprisingly, *Rbpms* and *Rbpms2* double CM-specific KO mice died before E13.5 and exhibited sarcomere disarray, whereas single KOs survive to adulthood albeit manifesting noncompaction (*Rbpms^cmKO^*) or without overt phenotype (*Rbpms2^cmKO^*). These findings suggest that RBPMS and RBPMS2 possess both distinct and overlapping functions in CMs. Although mis-splicing in cardiac genes can be seen in all single and double KOs, the splicing signature of double KO hearts shifts more markedly towards non-cardiac tissues, suggesting RBPMS proteins cooperatively promote cardiac splicing pattern while repressing non-cardiac splicing patterns across the genome. The splicing signature shift is partly contributed by the more prominent alterations in the splicing of sarcomere genes in double KO mice compared with single KO mice, which correlate with the severe phenotype in double KO embryos. Taking advantage of the well-defined RBPMS/RBPMS2 RNA binding motif, we found that the splicing regulation of RBPMS and RBPMS2 has the characteristics of “positional effects”, i.e., the binding location on pre-mRNA dictates whether they function as splicing activators or repressors; and the intrinsic features of the target exon, including CAC motif abundance and alternative 5’ splice site strength, likely determine the requirement for one or two RBPMS proteins for splicing regulation. Our study sheds light on the functional importance of cardiac SF cooperation in retaining cardiac splicing signature during heart development.

A recent report found that the splicing of several essential cardiac genes, including *Camk2d, Ryr2, Tpm1* and *Pdlim5*, are regulated by multiple cardiac SFs,^9^ but the functional importance of this cooperation remains largely unknown. In contrast, the collaboration among cardiac transcription factors (TFs) has been well documented. Cardiac TFs have been found to co-occupy at promoters or enhancers to regulate the transcription of cardiac genes.^24,57,58^ Moreover, the cooperation between cardiac TFs is functionally important, as *Nkx2-5* and *Tbx5* double KO embryos have far more severe phenotypes than *Nkx2-5* or *Tbx5* single KO embryos, due to the cooperative regulation of NKX2-5 and TBX5 on essential cardiac genes.^59^ Our study provides the first example of cooperation between cardiac splicing factors essential for heart development, as double KO of *Rbpms* or *Rbpms2* leads to embryonic lethality before E13.5 whereas single KO of *Rbpms* or *Rbpms2* survives to adulthood. Notably, the sarcomere disarray in double KO was not observed in single KOs, indicating RBPMS and RBPMS share a redundant yet indispensable role in regulating sarcomere assembly.

Splicing of genes essential for cardiac contractile function, e.g., sarcomere genes, calcium handling genes, is extensively regulated by cardiac SFs.^9^ However, the cardiac phenotypes of cardiac SF KO mice vary widely, with very few mouse models showing severe phenotypes like sarcomere disarray,^60,61^ suggesting that cardiac SFs may collaborate to ensure the proper splicing of sarcomere genes. This is evidenced by RBPMS and RBPMS2: despite the observation that ablating *Rbpms2* leads to sarcomere disarray in zebrafish,^20^ deleting *Rbpms* or *Rbpms2* in mice, as shown in our study and previous studies,^12,34^ does not. One feasible explanation is that RBPMS or RBPMS2 can compensate for each other’s function in regulating sarcomere assembly in murine heart, which is not possible in zebrafish heart that only expresses RBPMS2.^20^ To test the hypothesis that RBPMS and RBPMS2 can collectively regulate the splicing of sarcomere genes to support normal sarcomere assembly, we analyzed the splicing changes in the hearts of *Rbpms^cmKO^, Rbpms2^cmKO^*, and *Rbpms/Rbpms2^dcmKO^*, and identified alternative splicing (AS) events that are distinctly or commonly regulated by RBPMS proteins, most of which are of skipped exon (SE) or mutual exclusive exon (MXE) types. To gain insight into the functional relevance of these dysregulated AS events, we performed gene ontology (GO) analysis and found all three KOs enriched GO terms related to cardiac contractile apparatus. However, the splicing changes in genes essential cardiac contractile function are much more pronounced in double KO compared with single KOs, suggesting RBPMS proteins collectively contribute to the proper splicing of these genes. Next, we defined “cardiac splicing signature” by comparing splicing pattern of the heart to other organs, using RNA-seq datasets collected in various tissue types at different developmental time points.^46^ Interestingly, double KO samples tend to cluster with non-cardiac tissue samples while single KO samples largely stayed with “cardiac side”, indicating that RBPMS proteins collaboratively maintain the cardiac splicing signature in developing CMs.

Using the well-defined RBPMS/RBPMS2 RNA binding motif, we constructed the RNA splicing maps of RBPMS proteins by calculating motif enrichment around cassette exons promoted or repressed by RBPMS and/or RBPMS2. Our analysis revealed that RBPMS proteins tend to activate splicing when binding to the downstream intron close to the alternative 5’ss (Figure 7A). This “positional effects” of RBPMS splicing regulation resembles that of neural-specific SF Nova2, whose binding proximal to the alternative 5’ss also enhances splicing.^48,54^ Interestingly, binding to the alternative 5’ss appears to be the common mechanism for SFs to activate splicing, as similar behaviors have been observed with MBNL1,^62^ MBNL2,^63^ PTBP1,^52^ TIA,^55^ QKI,^53^ and RBFOX2.^7,64^ This activation effect is likely through enhancing U1 snRNP binding to the 5’ss and subsequent spliceosome assembly on the downstream intron^48,52,55^ (Figure 8A). Our analysis also suggested that RBPMS proteins tend to repress splicing when binding to the cassette exon itself and to the upstream constitutive 5’ss (Figure 7A), reminiscent of Nova, whose exonic binding blocks U1 snRNA binding at the downstream alternative 5’ss,^48^ and TIA, whose binding close to the constitutive 5’ss provides competition advantage over the alternative 5’ss^55^ (Figure 8B).

**Figure 8.**
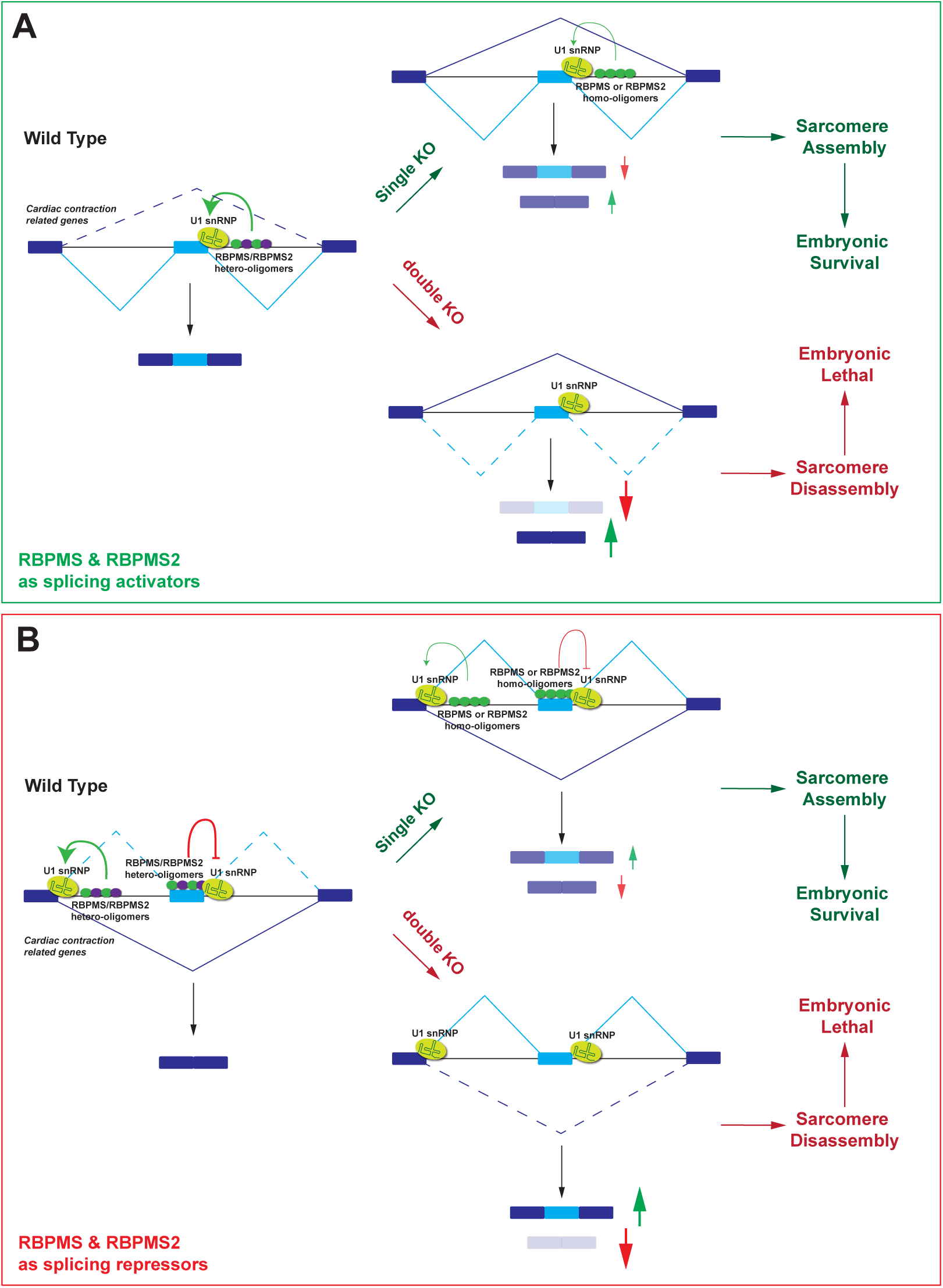
RBPMS/RBPMS2 hetero-oligomers function as splicing activators or repressors in CMs. **(A)** RBPMS and RBPMS2 act as splicing activators: RBPMS and RBPMS2 form hetero-oligomers and bind to CAC motifs at the alternative 5’ss of skipped exons (SEs), where they facilitate U1 snRNP binding to the 5’ss, subsequent spliceosome assembly on the downstream intron, and exon inclusion. In *Rbpms* or *Rbpms2* single KO, the remaining RBPMS member retains partial ability of RBPMS/RBPMS2 hetero-oligomers to promote the inclusion of exons essential for sarcomere assembly. As a result, single KO mice’ sarcomeres assemble normally and survive to adulthood. However, losing both RBPMS and RBPMS2 proteins in double KO leads to severe mis-splicing of contraction-related genes, sarcomere disassembly and embryonic lethality. **(B)** RBPMS and RBPMS2 act as splicing repressors: RBPMS/RBPMS2 hetero-oligomers bind to the skipped exon itself and inhibit the recruitment of U1 snRNP to alternative 5’ss. They may also occupy the constitutive 5’ss and provide competition advantage over the alternative 5’ss. Both scenarios lead to repressed splicing of the regulated exon and favor its skipping. While mis-splicing of contraction-related genes (inclusion of inappropriate exons in this case) in single KOs is not severe enough to cause sarcomere disarray and embryonic lethality, deleting both *Rbpms and Rbpms2* in mice is clearly incompatible with life.

Another intriguing aspect of RBPMS and RBPMS2 is the requirement for one or two proteins for splicing regulation. To investigate this, we compared the characteristics of three sets of exons: exons only regulated by RBPMS, exons only regulated by RBPMS2, and exons regulated by both. We found that exons commonly promoted by RBPMS proteins tend to have higher CAC motif coverage at their alternative 5’ss (Figure 7B). A recent report discovered that RBPMS undergoes homomeric oligomerization through its C-terminal intrinsically disordered regions (IDRs), which allows RBPMS homo-oligomer to bind multiple CAC motifs on the pre-mRNA and interact with several splicing co-regulators.^65^ Although RBPMS and RBPMS2 contain almost identical RNA recognition motifs (RRMs),^14^ the differences in their amino acid sequence are larger in other regions (Supplemental Figure 6A), which may enable them to interact with different binding partners. Thus, a higher concentration of CAC motifs may have a better chance to recruit both RBPMS and RBPMS2, which in turn provides access to a more diverse pool of splicing co-regulators that could help to stabilize the splicing regulatory complexes.

Interestingly, RBPMS is found to physically interact with RBPMS2 in CMs,^34^ suggesting they may not only bind to the same set of CAC motifs but also form hetero-oligomers. Notably, most genes important for cardiac contractile function require both RBPMS and RBPMS2 for splicing regulation (Figure 5D), indicating they may utilize them as “double safety” to ensure their proper splicing. In addition, these genes tend to have stronger alternative 5’ss (Figure 7C), which could be another layer of insurance for efficient splicing. Future studies are warranted to test the impact of disrupting the formation of RBPMS-RBPMS2 hetero-oligomers on splicing regulation.

Lastly, the functional interplay among splicing factors within the same family seems to be a common phenomenon in many tissue types: neural-specific splicing factors Nova1 and Nova2 recognize highly similar RNA motifs^66^ and can regulate the same set of exons;^67^ and broadly expressed MBNL1 and MBNL2 are interchangeable in splicing regulation.^62^ However, Nova1 and Nova2 show reciprocal expression patterns in the brain, and MBNL1 and MBNL2 are differentially required in brain and skeletal muscles due to their variable expression levels in those tissues.^63^ Whether RBPMS and RBPMS2 exhibit distinct expression patterns in the heart remains unclear, as all tested RBPMS and RBPMS2 antibodies recognize both proteins (Supplemental Figure 1C-D and data not shown), making them unsuitable for immunofluorescence analysis. However, based on single-cell transcriptomic data across multiple human tissues available in the GTEx Portal database, we observe widespread expression divergence of these paralogous splicing factors across different tissue and cell types (Supplemental Figure 6B). Despite considerable sequence and expression divergence, RBPMS and RBPMS2 have retained complementary roles as splicing factors integral to heart development, highlighting the specificity and highly refined nature of alternative splicing regulation in cardiomyocytes.

## Non-standard Abbreviations and Acronyms

RBPMS: RNA Binding Protein with Multiple Splicing
RBPMS2: RNA Binding Protein with Multiple Splicing 2
CM: cardiomyocyte
DEG: differentially expressed genes
AS: precursor messenger RNA (pre-mRNA) alternative splicing
SF: splicing factor
SE: skipped exon
MXE: mutual exclusive exon

## Acknowledgements

We are grateful to Kristen Jepsen (UCSD IGM Genomics Center) and Jennifer Santini (UCSD Microscopic Core Facility, supported by NIH grant P30NS047101) for their technical assistance.

## Sources of Funding

T. W. is supported by startup funding from the Masonic Medical Research Institute. J.C. is funded by grants from the National Heart, Lung, and Blood Institute and J.C. holds an American Heart Association Endowed Chair in Cardiovascular Research.

## Disclosures

None.

## Supplemental Materials

Supplemental Figures I - VI

Supplemental Excel Files I - II

---

**Supplemental Table 1. Gene ontology analysis of upregulated or downregulated differentially expressed genes (DEGs) in *Rbpms^cmKO^, Rbpms2^cmKO^*, and *Rbpms/Rbpms2^dcmKO^*hearts.**

**Supplemental Table 2. Gene ontology analysis of dysregulated skipped exon (SE) and mutual exclusive exon (MXE) splicing events in *Rbpms^cmKO^, Rbpms2^cmKO^*, and *Rbpms/Rbpms2^dcmKO^*hearts.**

## Notes

### Competing Interest Statement

The authors have declared no competing interest.

### Summary of Updates

This revision is to add supplemental figures to the manuscript.

