## Supplemental Figures 1-6 for "RBPMS and RBPMS2 Cooperate to Safeguard Cardiac Splicing"

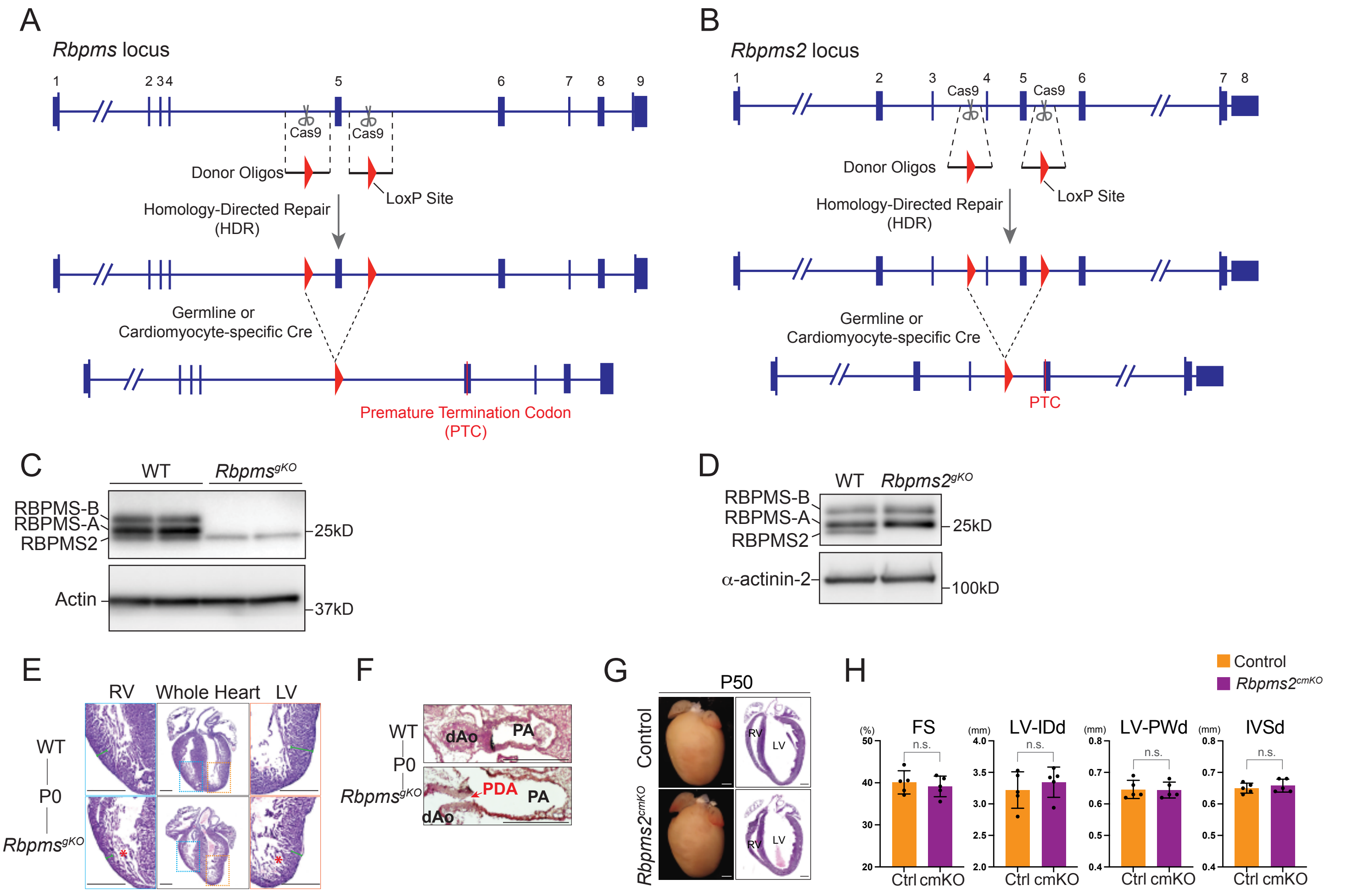


**Supplemental Figure 1. Generation of *Rbpms* and *Rbpms2* floxed mice using CRISPR-Cas9.** **(A)** The scheme of inserting 2 LoxP sites flanking exon 5 of *Rbpms* gene, whose deletion by Cre will introduce a premature termination codon (PTC) and lead to nonsense-mediated RNA decay (NMD) of *Rbpms* mRNA. **(B)** The scheme of inserting 2 LoxP sites flanking exon 4 and 5 of *Rbpms2* gene, which enables their deletion by germline or CM-specific Cre, introducing a PTC and leading to NMD of *Rbpms2* mRNA. **(C)** Pan-RBPMS western blot analysis shows RBPMS-A and RBPMS-B are completely absent in *Rbpms*$^{gKO}$ hearts at E13.5, whereas RBPMS2 is unaffected. The shorter RBPMS isoform, RBPMS-A, was expressed in higher level than the longer RBPMS-B and RBPMS2 in the heart. Actin is used as a loading control. **(D)** Pan-RBPMS western blot analysis shows RBPMS2 is completely abolished in *Rbpms2*$^{gKO}$ heart at E17.5, while RBPMS-A and RBPMS-B are unaffected. a-actinin-2 is used as a loading control. **(E)** H&E-stained whole heart sections from wildtype and *Rbpms*$^{gKO}$ mice at P0. Red asterisks indicate noncompaction in *Rbpms*$^{gKO}$ mice. Green rulers indicate compact layer thickness. Scale bar: 0.5 mm. **(F)** H&E staining showed the presence of patent ductus arteriosus (PDA) in *Rbpms*$^{gKO}$ mice. dAo: descending aorta, PA: pulmonary artery. Scale bar: 0.5 mm. **(G)** Wholemount images (left) and H&E-stained sections (right) from control and *Rbpms2*$^{cmKO}$ mice at P50. LV, left ventricle; RV, right ventricle; Scale bar: 1 mm. **(H)** Echocardiography analysis of control (n=5) and *Rbpms2*$^{cmKO}$ (n=5) mice at P50. n.s., not significant (Welch's t-test). FS: fraction shortening; LV-IDd: LV internal dimension, diastolic; LV-PWd: LV posterior wall thickness, diastolic; IVSd: interventricular septum thickness, diastolic.

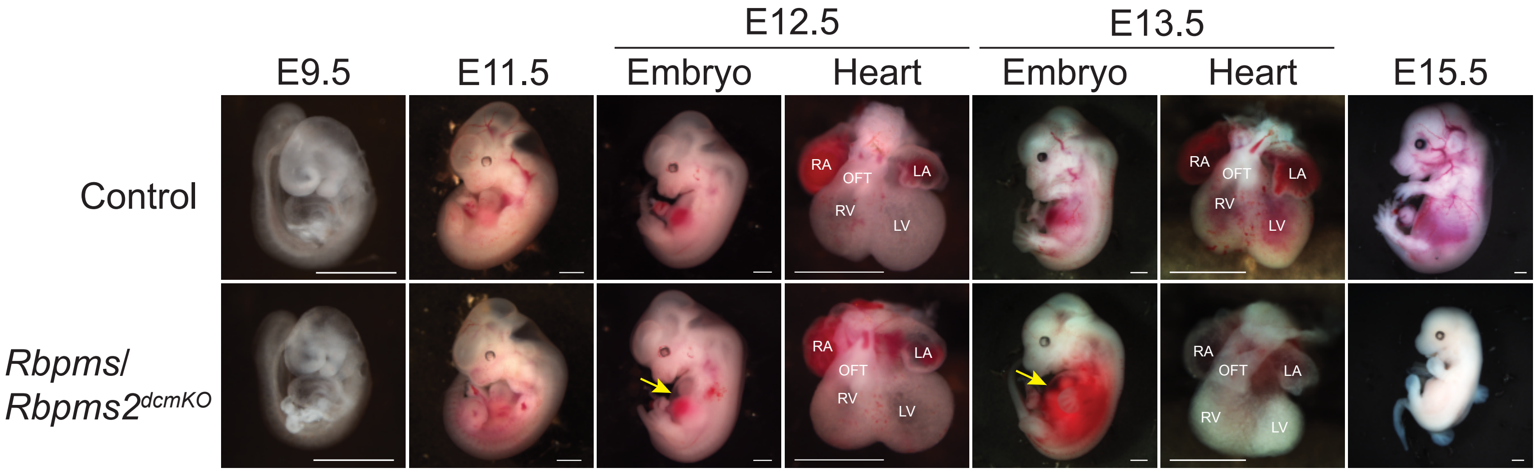


**Supplemental Figure 2. Gross morphology of *Rbpms/Rbpms2* CM-specific double KO (*Rbpms/Rbpms2*$^{dcmKO}$) embryos.** Representative brightfield images of *Rbpms/Rbpms2*$^{dcmKO}$ and littermate control embryos or embryonic hearts from E9.5 to E15.5. Yellow arrows indicate pericardial effusion, indicative of insufficient cardiac function in embryos. Scale bar: 1 mm.

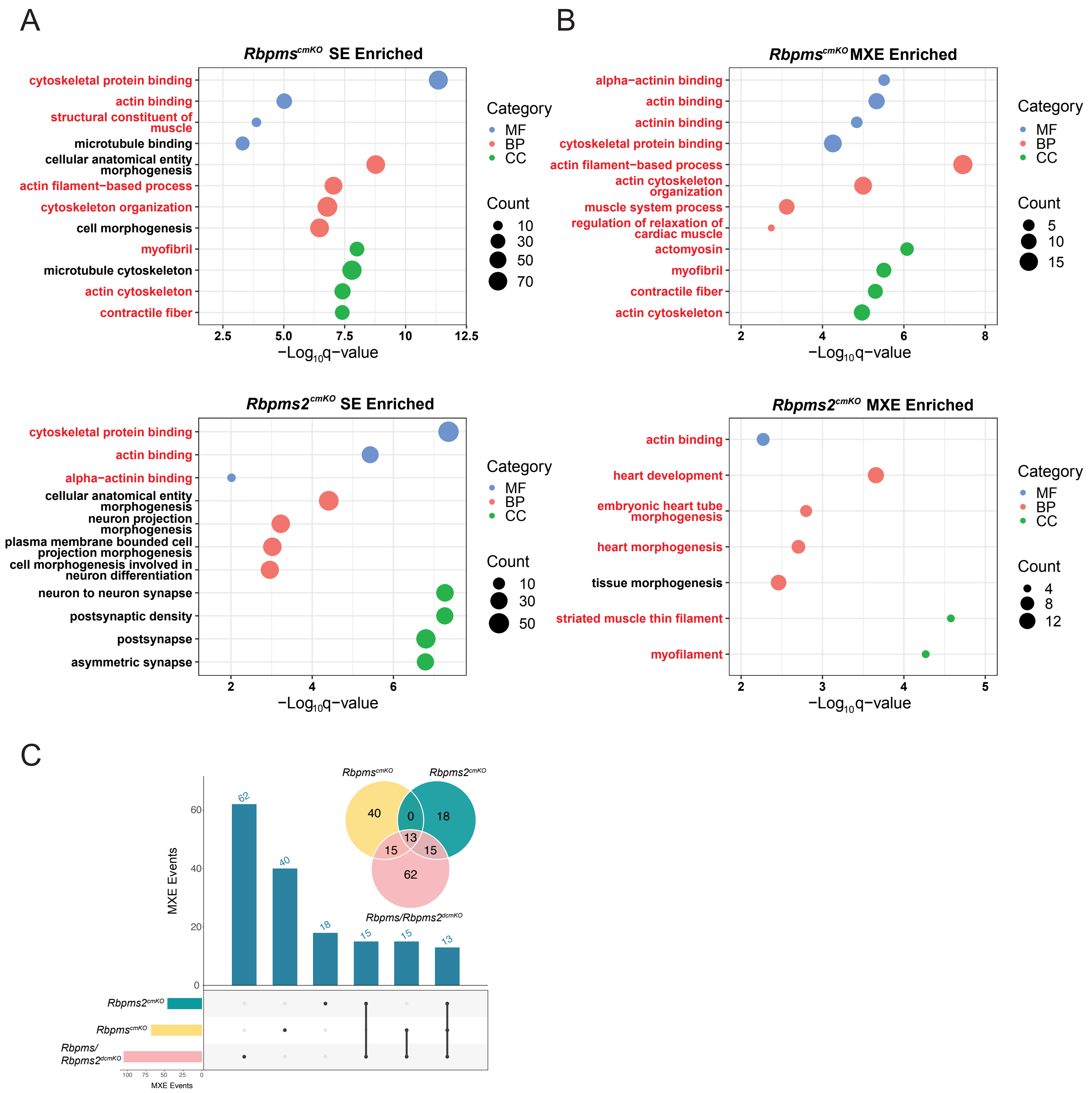


**Supplemental Figure 3. RBPMS and RBPMS2 cooperate in cardiac splicing. (A)** Gene ontology (GO) analysis of mis-spliced cassette/skipped exon (SE) events in *Rbpms*$^{cmKO}$ and *Rbpms2*$^{cmKO}$ hearts. GO terms related to cardiac contraction are highlighted in red. BP, biological process; CC, cellular component; MF, molecular function. **(B)** GO analysis of mis-spliced mutual exclusive exon (MXE) events in *Rbpms*$^{cmKO}$ and *Rbpms2*$^{cmKO}$ hearts. GO terms related to cardiac contraction are highlighted in red. **(C)** Overlap of mis-spliced MXE events among *Rbpms*$^{cmKO}$, *Rbpms2*$^{cmKO}$ and *Rbpms/Rbpms2*$^{dcmKO}$ hearts.

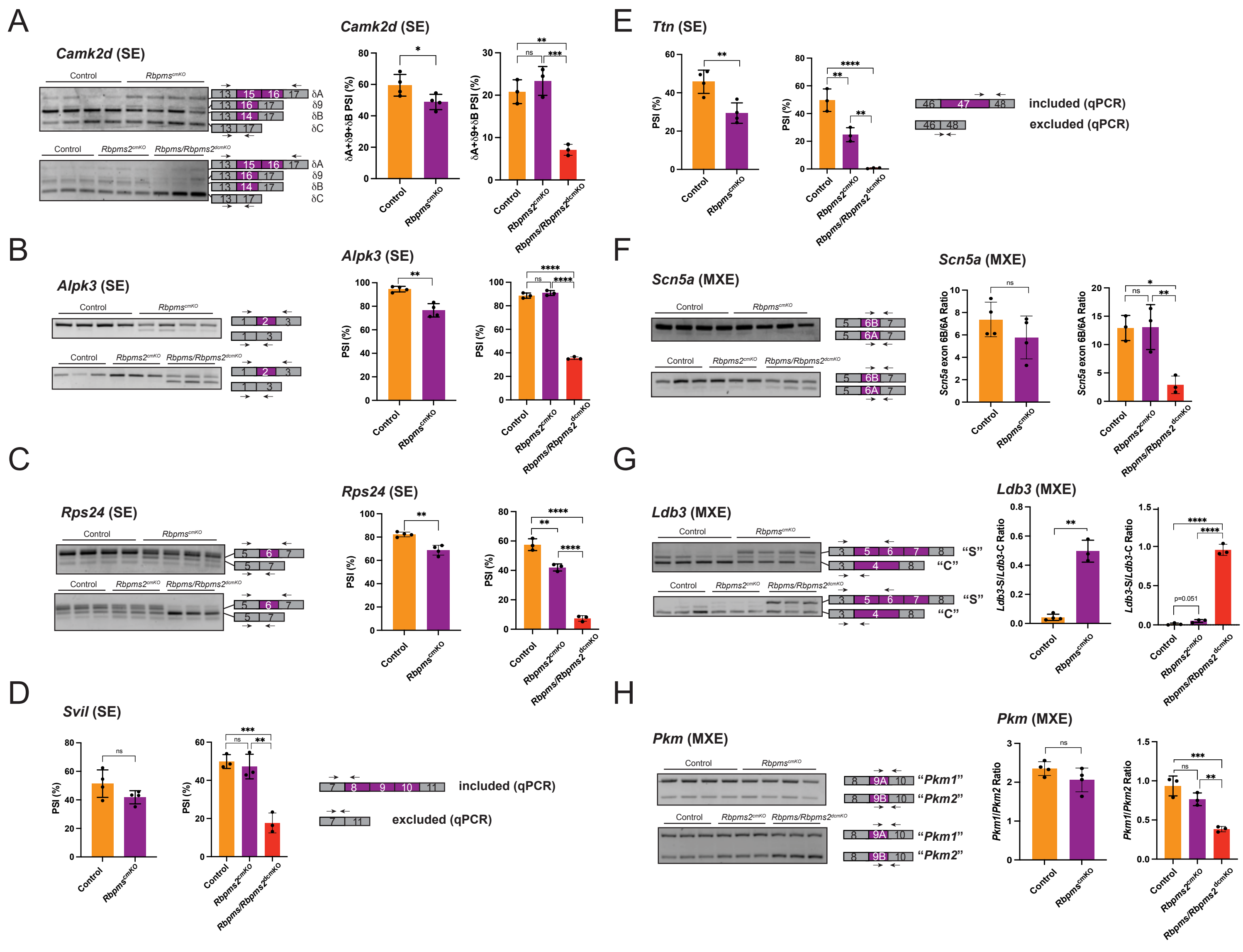

**Supplemental Figure 4. RBPMS and RBPMS2 cooperate in cardiac splicing (continued).**
**(A-H)** RT-PCR validation of mis-spliced skipped exon (SE) or mutual exclusive exon (MXE) alternative splicing (AS) events using RNA samples from samples from *Rbpms*$^{cmKO}$, *Rbpms2*$^{cmKO}$ and *Rbpms/Rbpms2*$^{dcmKO}$ hearts. *p<0.05, **p<0.01, ***p<0.001, ****p<0.0001. Statistical significance was determined by Welch's t-test (two groups) or one-way ANOVA (three groups).

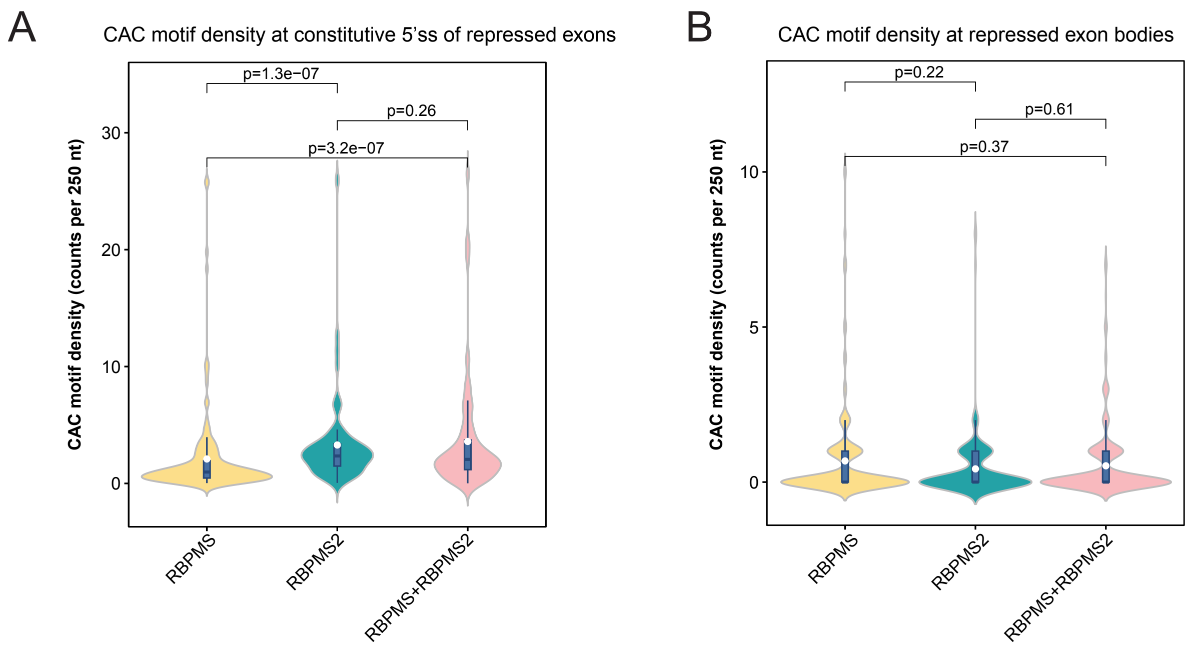


**Supplemental Figure 5. RNA maps of RBPMS and RBPMS2 in CMs.** Comparison of RBPMS/RBPMS2 motif coverage at constitutive 5'ss **(A)** and within **(B)** SE exons repressed by RBPMS proteins individually or together. Statistical significance is determined by pairwise Wilcoxon rank-sum tests. Benjamini-Hochberg method was applied to the obtained adjusted p-values due to multiple comparisons.

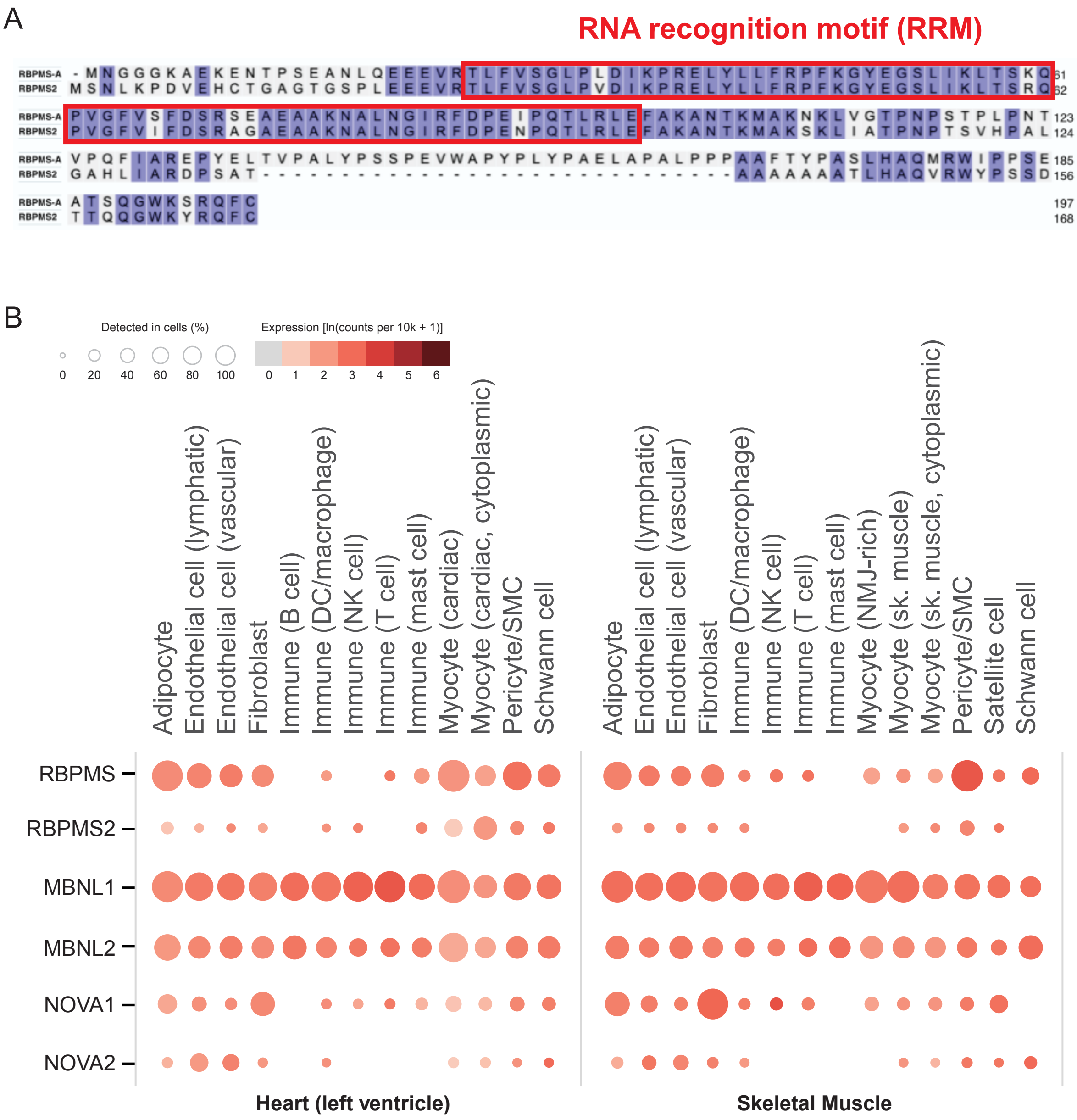


**Supplemental Figure 6. Cooperation of paralogous splicing factors. (A)** Alignment of murine RBPMS (RBPMS-A isoform) and RBPMS2 protein sequences. RNA recognition motifs (RRMs) are indicated by red boxes. **(B)** Expression pattern of paralogous splicing factors in different cell types in human heart (left ventricle) and skeletal muscle (gastrocnemius). Data were derived from single-cell gene expression from the GTEx Multi-Gene Single Cell Query, accessed via the GTEx Portal (GTEx Analysis Release V8, dbGaP Accession phs000424.v8.p2).
